# Assessing Goodness-of-Fit in Marked-Point Process Models of Neural Population Coding via Time and Rate Rescaling

**DOI:** 10.1101/2020.01.24.919050

**Authors:** Ali Yousefi, Yalda Amidi, Behzad Nazari, Uri. T. Eden

## Abstract

Marked-point process models have recently been used to capture the coding properties of neural populations from multi-unit electrophysiological recordings without spike sorting. These ‘clusterless’ models have been shown in some instances to better describe the firing properties of neural populations than collections of receptive field models for sorted neurons and to lead to better decoding results. To assess their quality, we previously proposed a goodness-of-fit technique for marked-point process models based on time-rescaling, which for a correct model, produces a set of uniform samples over a random region of space. However, assessing uniformity over such a region can be challenging, especially in high dimensions. Here, we propose a set of new transformations both in time and in the space of spike waveform features, which generate events that are uniformly distributed in the new mark and time spaces. These transformations are scalable to multi-dimensional mark spaces and provide uniformly distributed samples in hypercubes, which are well suited for uniformity tests. We discuss properties of these transformations and demonstrate aspects of model fit captured by each transformation. We also compare multiple uniformity tests to determine their power to identify lack-of-fit in the rescaled data. We demonstrate an application of these transformations and uniformity tests in a simulation study. Proofs for each transformation are provided in the Appendix section. We have made the MATLAB code used for the analyses in this paper publicly available through our Github repository at https://github.com/YousefiLab/Marked-PointProcess-Goodness-of-Fit

## 1 Introduction

In recent years, marked point process models have become increasingly common in the analysis of population neural spiking activity [1–3]. For multi-unit spike data, these models directly relate the occurrences of spikes with particular waveform features to the biological, behavioral, or cognitive variables encoded by the population, without the need for a separate spike-sorting step. For this reason, these are sometimes called ‘clusterless’ neural models. Clusterless models have been shown to capture coding properties for spikes that cannot be sorted with confidence and to lead to improved population decoding results in place field data from rat hippocampus during spatial navigation tasks [1–3]. Additionally, avoiding a computationally intensive spike-sorting step allows for neural decoding to be implemented in real-time, closed-loop experiments.

A critical element of any statistical modeling procedure is the ability to assess the goodness-of-fit between a fitted model and the data. For point process models of sorted spike train data, effective goodness-of-fit methods have been developed based on the time-rescaling theorem [4, 5]. Previously, we developed an extension of the time-rescaling theorem for marked point processes, which given the correct model, rescales the observed spike and mark data to a uniform distribution in a random subset of a space of the marks and rescaled times [6]. We can then use established statistical tests for uniformity to assess whether the model used for rescaling is consistent with the observed data. However, several challenges still limit the efficient application of these methods to marked point process models, in some cases. For models with high-dimensional marks representing the waveform features, computing the space in which the rescaled data should be uniform can be computationally expensive [6]. Since this space is random and typically not convex, the number of statistical tests for uniformity is limited to those that can be applied in general spaces. Finally, of the multitude of uniformity tests, it is often not clear which should be applied to the rescaled data.

Here, we propose several extensions to this goodness-of-fit approach based on combinations of time and mark scaling, which for a correct model, transform the observed spike and waveform data to uniformly distributed samples in a hypercube. This in turn, simplifies and opens up more options for assessing uniformity. We discuss properties of each transformation and demonstrate which aspects of model lack-of-fit are better captured using each. Finally, we perform a simulation analysis to compare and contrast the transformations proposed here – along with the multiple uniformity tests - to assess different models’ fit to the simulated data.

Our goal here is not to identify one single, best transformation and uniformity test for assessing goodness-of-fit of marked point process models; instead, we aim to provide a toolbox of methods to identify multiple ways in which a model may fail to capture structure in the data and to provide guidance about which methods are most likely to be useful in different situations. We also developed an interactive and easy-to-use toolbox for the transformations and uniformity tests described here, to assist other researchers in applying these goodness-of-fit techniques in their analysis of neural spike trains.

The paper is organized as follows: we first introduce each transformation in detail and briefly discuss their core properties. We then discuss different uniformity tests and their main attributes. We then go through a simulation example and compare goodness-of-fit results for the true and a set of alternative generative models. We finish the paper with theoretical proofs that the transformations under the correct model yield uniform samples.

## 2 Marked-Point Process to Uniform Transformation

In this section, we introduce two transformations that take a dataset of spike times and waveforms from a marked point process model to a set of identically distributed uniform samples on the hypercube [0 1]^*d*+1^, where *d* is the dimension of the mark used to describe the spike waveform features in the model. We also discuss properties of both transformations and explain which features of model misfit can be better captured by each transformation.

A marked point process model is defined by a joint mark intensity function (JMIF), 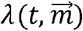 where *t* represents time, and 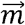 represents a vector mark describing spike waveform features. 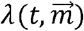 is defined so that the likelihood of observing a spike at time *t* with a waveform with features in a neighborhood 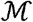 of 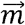 is given by,

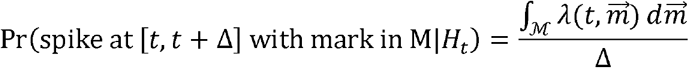

A marked point process model expresses this intensity as a function of any signals or covariates encoded by the neural population, *x*(*t*), and the history of spikes and waveforms up to time *t*, *H*_*t*_. Using this joint mark intensity, we can compute the ground intensity 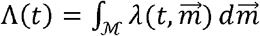, where 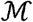 is the full space of marks, which defines the intensity of observing any spike at time *t*. Similarly, we can define the mark intensity 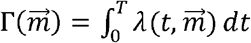.

For an observation interval, [0 *T*], we observe a sequence of spike events at times 0 = *s*_0_ < *s*_1_ < *s*_2_ < ··· < *s*_*i*_ < ··· < *s*_*N*(*T*)_ < *T* with associated marks 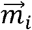, for *i* = 1, …, *N*(*T*) with joint mark intensity function 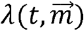. We assume this joint mark intensity function is integrable over both time and mark space. The notation we use to define the data and model components are listed in Table 1.

**Table 1:**
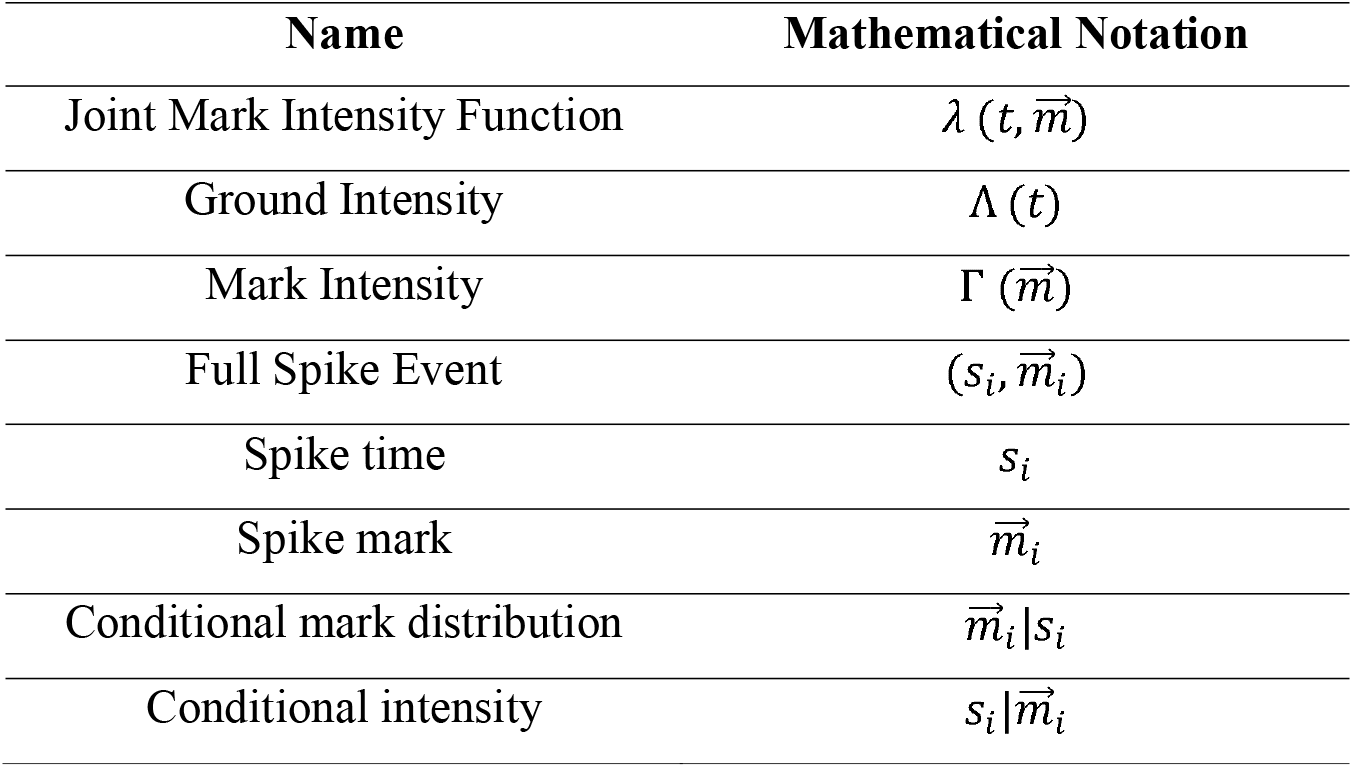
Notation for the marked point process model and data

In the following subsections, we present the transformations and associated uniformity tests.

### 2.1 Interval-Rescaling Conditional Mark Distribution Transform (IRCM)

This algorithm requires the computation of the ground intensity – Λ(*t*) – followed by the conditional mark distribution 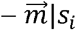. We first rescale the inter-spike intervals across all observed spikes based on the ground intensity. We then rescale each mark dimension sequentially, using a Rosenblatt transformation [7] based on the conditional mark distribution given the spike time. The order of conditioning for the mark features can be specified directly or selected randomly. The dimension of new data samples is *d* + 1, where *d* is the dimension of mark space. The transformed data samples are i.i.d with a uniform distribution in the hypercube [0 1]^*d*+1^. The following table presents the first algorithm, which we call the Interval-Rescaling Conditional Mark Distribution Transform or IRCM.

#### Algorithm 1

Interval-Rescaling Conditional Mark Distribution Transform (IRCM)

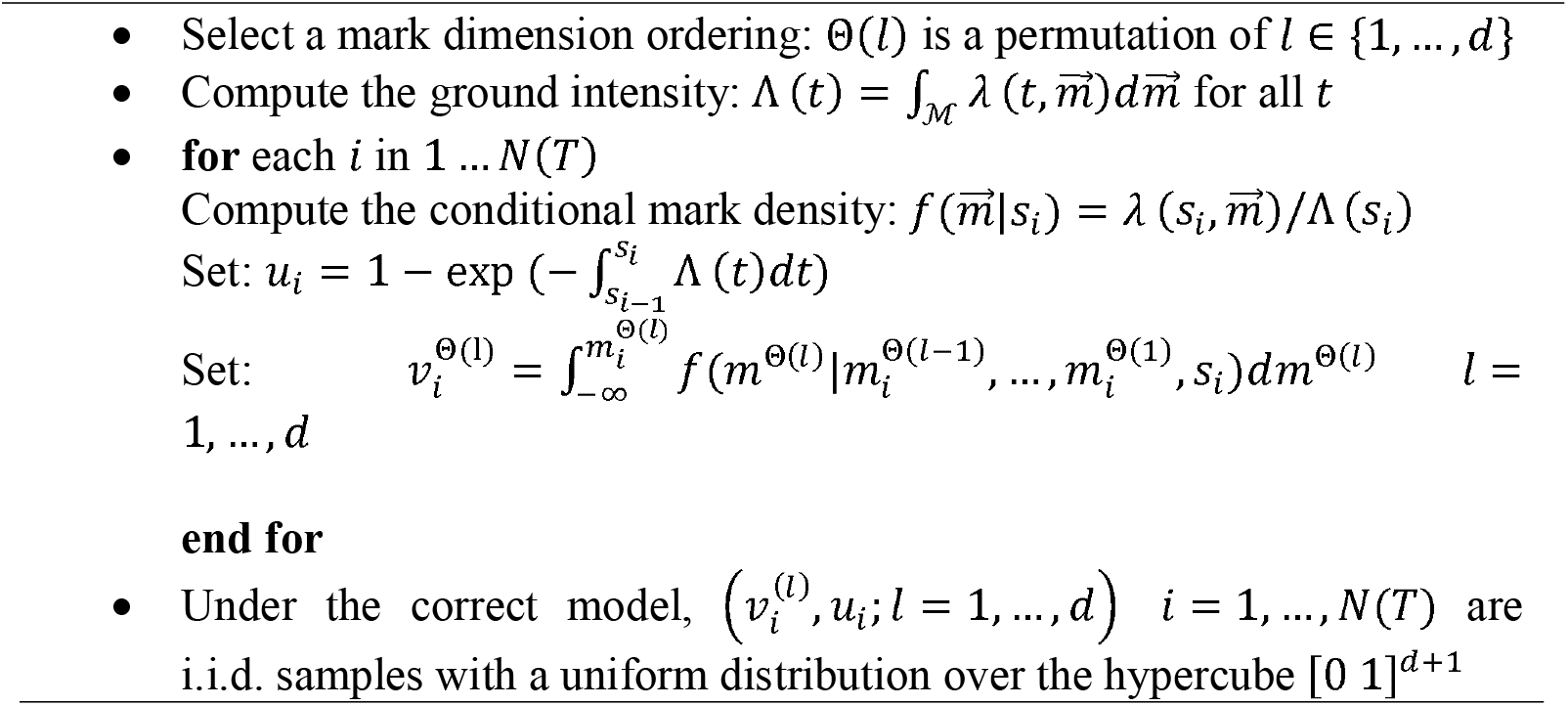

No matter which ordering we select for the mark components, the i.i.d. and uniformity properties will hold for the true model. The theoretical proof of IRCM transformation is included in Appendix A.1 section - Theorem 1 and Corollaries 1, and 2.

The rescaled data samples from the IRCM transformation not only provide insight about the overall quality of the proposed joint mark intensity function, but also reveal finer aspects of the model fit or misfit. The *u*_*i*_ samples are computed using only the spike times and the estimated ground intensity model and can be used separately to assess the goodness-of-fit of the temporal component of the model to the unmarked spike times using the time-rescaling theorem [5]. The 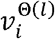 samples for a fixed dimension, *l*, are computed using only marks Θ(1),…,Θ(*l*) and the conditional mark density, *f*(*m*^Θ(*l*)^|*m*^Θ(*l−1*)^,…,*m*^Θ(*1*)^, *s*_*i*_); if samples 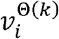 for *k* < *l* are uniform but 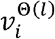 are not, this suggests specific lack of fit in modeling the coding properties of the waveform features associated with mark dimension *l*.

### 2.2 Mark Density Conditional Intensity Transform (MDCI)

This algorithm requires rescaling time separately for each spike, based on its joint mark intensity. This can potentially break the ordering of spikes with different waveform features, while spikes with similar waveforms will tend to maintain their relative ordering. Next, the algorithm sequentially rescales each mark dimension, again based on a Rosenblatt transformation [7]. Like IRCM, we can choose any ordering for the mark features, or select a random ordering. Distinct from IRCM, this transformation does not depend on the time of the spike, only on its mark value. The table below describes this mapping, which is called Mark Density Conditional Intensity Transform or MDCI.

#### Algorithm 2

Mark Density Conditional Intensity Transform (MDCI)

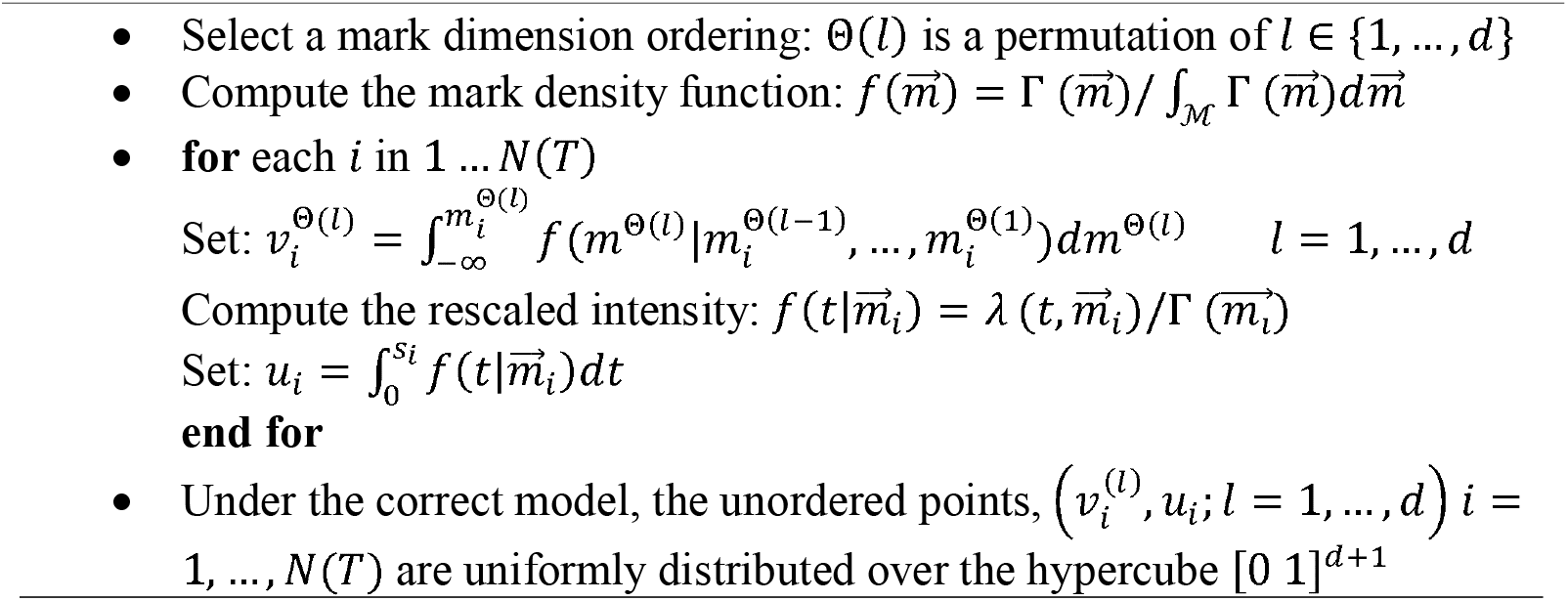

The proof that the MDCI transformation under the true conditional intensity model leads to uniform samples is included in Appendix A.2 section – Theorem 2.

The key difference between the IRCM and MDCI transforms is that the IRCM transforms the inter-spike intervals independent of their marks and then transforms each mark based on the intensity of spikes with that waveform at the observed time, while the MDCI transforms the marks independent of when the corresponding spikes occur and then transforms time differently for each spike waveform. For neural spiking models, the IRCM examines the intervals between spikes, and tends to mix the marks so that spikes with similar waveforms may end up far apart in the transformed mark space; inversely, the MDCI tends to leave spikes with similar waveforms nearby in the transformed mark space, while mixing up the spike timing from different neurons. Another important difference is that, for the correct model, the IRCM generates i.i.d. uniform samples while the MDCI samples are not independent. However, the set of all the unordered MDCI samples do have a joint uniform distribution. We therefore expect these transforms to allow us to determine separate aspects of lack of fit. The misfit associated with the model of individual neurons or particular waveform features might be better assessed using MDCI while misfit associated with interactions between neurons might be better assessed using IRCM. We investigate these expectations in the Simulation section below.

In this section, we described two algorithms which take marked point process data and map them to uniformly distributed samples in a hypercube, [0 1]^*d*+1^ based on their joint mark intensity. These methods allow for marks of arbitrary dimension. In Appendix B we describe one additional transformation, which applies in the specific case where the mark is scalar.

## 3 Uniformity Tests

There are a multitude of established uniformity tests for one-dimensional data; however, the number of established, robust, multi-dimensional uniformity tests is more limited. Pearson’s chi-square test can be used to assess uniformity by partitioning the space into discrete components and computing the number of samples in each [8, 9]. Another approach is to apply a multivariate Kolmogorov-Smirnov (KS) test [10], which uses a statistic based on the maximum deviation between the empirical multivariate cdf and that of the uniform to build a distribution free test for multi-dimensional samples. Other test statistics are derived from number-theoretic or quasi-Monte Carlo methods for measuring the discrepancy of points in [0 1]^*d*^ [11, 12]. Using Monte Carlo simulation, it is known that the finite-sample distribution of these statistics can be well approximated by a standard normal distribution [11, 12]. Two other approaches to assessing multivariate uniformity are based on distances between samples and the boundary of the hypercube [13] and distances between nearest samples, which leads to the computation of Ripley’s K function [14–16]. Fan [17] describes a test based on the *L*_2_ distance between the kernel density estimate of the underlying probability density and the uniform distribution. Other tests include those built upon order statistics [18], Friedman-Rafsky’s minimal spanning tree [19], or a weighted *K*-function [20–22]. There are several other multivariate uniformity tests which are not presented here; a comprehensive discussion of scalar and multivariate uniformity tests can be found in [23]. There are also uniformity tests specifically designed for two- and three-dimensional spaces including complete spatial randomness or bivariate Cramer-von Mises tests that are described in [24–26].

Here, we investigate a few of these approaches in terms of their ability to detect model misfit in rescaled samples from the spike transformations described above; the tests are a Pearson’s chi-square test [8], a multivariate KS test [10], the distance-to-boundary method [13], a discrepancy-based test [11], a test based on Ripley’s *K*-function [14–16], and a test using minimum spanning trees (MST) [27, 28]. The tests are described in detail in the cited literature and are expressed algorithmically in Table 2. These tests tend to be straightforward to implement with a few exceptions; Ripley’s *K*-function becomes computationally expensive to test in more than two dimensions; Pearson’s chi-square requires defining a set of sub-regions of the hypercube. The remaining tests do not require any parameters to be selected except for the test significance level.

**Table 2:** Uniformity Tests; *d* is the dimension of the data, *n* is the number of data samples, and *α* is the significance level

| Test Name | Method |
| --- | --- |
| **Pearson $\chi^2$ test [8]** | 1. Define $M$ sub-regions, $R_j$ $j = 1, \dots, M$ in the hypercube $R$<br>2. Let $p_j = \vert R_j\vert /\vert R\vert $, where $\vert R\vert $ represents the volume of region $R$.<br>3. Calculate $\chi^2 = \sum_{j=1}^M (r_j - np_j)^2 / np_j$ using the rescaled data samples, where $r_j$ is the number of samples in $R_j$<br>4. If $\chi^2 > \chi_{M-1}^2(1 - \alpha)$ reject the null hypothesis, where $\chi_{M-1}^2(1 - \alpha)$ is the inverse of the chi-square cdf with $M - 1$ degrees of freedom. |
| **Multivariate Kolmogorov-Smirnov test [10]** | 1. Define $D^{KS} = \sup_n \vert F(v_n^{(1)}, \dots, v_n^{(d)}, u_n) - u_n \prod_{i=1}^d v_n^{(d)}\vert $, where $F(\cdot)$ is the empirical multivariate CDF<br>2. Use Monte Carlo simulation to find the critical value $-C_\alpha$ for the test at the significance level of $\alpha$<br>3. Calculate $D^{KS}$ using transformed data samples.<br>4. If $D^{KS} > C_\alpha$ reject the null hypothesis |
| **Distance to boundary test [13]** | 1. Define $D(u, \delta S) = \min_{x \in \delta S} \ x - u\$, where $\delta S$ is the boundary of the hypercube and $\ \cdot\$ is Euclidean norm on $R^d$<br>2. Calculate $y_i = D(u_i, \delta S) / \max(D(u_j, \delta S))$ for $j = 1, \dots, n$ samples<br>3. Calculate $D^{KS} = \sup_i F(y_i) - 1 + (1 + y_i)^d$, where $F(\cdot)$ is the empirical CDF<br>4. If $D^{KS} > C_\alpha / \sqrt{n}$, where $C_\alpha$ is the KS critical value, reject the null hypothesis |
| **Discrepancy test [11]** | 1. Calculate $A_n = \sqrt{n}[(U_1 - M^d) + 2(U_2 - M^d)] / (\sqrt{5\xi_1})$,<br>-Appendix C.1 provides the definitions for $M, \xi_1, U_1$ and $U_2$<br>Under the null hypothesis, $A_n$ has a standard normal distribution.<br>2. If $\vert A_n\vert > z_{\alpha/2}$, reject the null hypothesis, where $z_{\alpha/2}$ is the critical value of a standard normal at significance level $\alpha$. |
| **Ripley statistics test [15, 16]** | 1. Compute the distance from each rescaled point to its nearest neighbor, $\vec{r} = \{r_1, \dots, r_s\}$<br>2. For each $r_k$ calculate Ripley's $K$-function statistics by $\hat{K}(r_k) = \sum_{u_i \neq u_j} I\{d(u_i, u_j) \leq r_k\}$<br>3. Let $\hat{K} = (\hat{K}(r_1) - B(r_1), \dots, \hat{K}(r_s) - B(r_s))$ where $B(r_k) = -\frac{8}{3}r_k^3 + \frac{1}{2}r_k^4$<br>4. Calculate $T^2 = \left\ \Sigma^{-\frac{1}{2}}(\hat{K} - \pi \vec{r}^2) \right\ ^2$ - under the null hypothesis, $T^2$ has a chi-square distribution with $d$ degree of freedom -Appendix C.2 provides the definition for $\Sigma$.<br>5. If $T^2 > \chi_d^2(\alpha)$, where $\chi_d^2(\alpha)$ is the chi-square critical value, reject the null hypothesis |
| **Minimal Spanning Tree (MST) test [27, 28]** | 1. Draw $m$ multi-variate uniform sample points<br>2. Calculate the number of degree pairs, $C$, in the MST that share a common node.<br>3. If $d_i$ is the degree of $i^{th}$ node, $C$ is defined by $C = 1/2 \sum_{i=1}^N d_i(d_i - 1)$, $N = m + n$<br>4. Calculate the number of edges linking a point from the generated data to uniform sample points $-T$<br>5. Under the null hypothesis assumption, calculate $\text{Var}[T\vert C] = \frac{2mn}{N(N-1)} \left( \frac{2mn-N}{N} + \frac{C-N+2}{(N-2)(N-3)} [N(N-1) - 4mn + 2] \right)$ and $E[T] = \frac{2mn}{N+1}$<br>6. Calculate $D = (T - E[T]) / \sqrt{\text{Var}[T\vert C]}$<br>7. If $D < z_\alpha$, reject the null hypothesis, where $z_\alpha$ is the α-quantile of the standard normal distribution |

The data transformations require the selection of an ordering of the mark dimensions; the uniformity tests can be applied to one particular ordering or can be modified to allow for assessment across multiple permutations of orderings. In such cases, the test procedures should be adjusted for multiple comparisons [29].

## 4 Simulation Study

In this section, we demonstrate an application of the IRCM and MDCI transformations along with the multiple uniformity tests described in the Table 2 to assess their ability to measure goodness-of-fit in simulation data. We first describe how the simulation data is generated, and then examine the transformations and goodness-of-fit results.

### 4.1 Simulation Data

We generate simulated spiking data using a marked-point process intensity model consisting of two connected neurons encoding a simulated position variable, x_*t*_. *x*_*t*_ is modeled as an AR1 process,

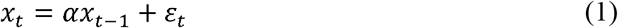

where *α* is set at 0.98 and *ε*_*t*_ is a zero mean white noise process with a standard deviation of 0.3. The neurons’ spiking activity depends on *x*_*t*_ and on previous spiking; each neuron has a refractory period and neuron 2 has an excitatory influence on neuron 1. We generate the simulated spike data in discrete time using a step size of 1 millisecond, based on the following joint mark intensity function

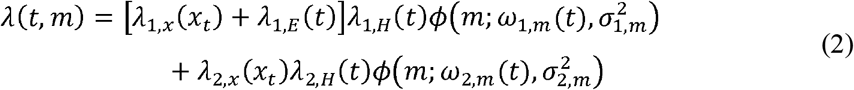

where *ϕ*(*x*;*w*,*σ*^2^) is the pdf of a normal distribution with mean w and variance *σ*^2^ at point *x*, used to represent the variability of the spike waveform marks. In Equation (2)

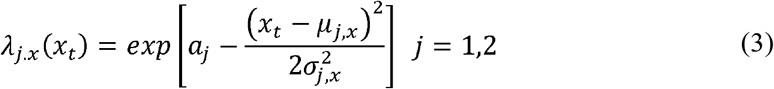

represents the receptive fields of neuron 1 and 2, *μ*_*j,x*_ and 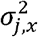 define the field center and width, and *a*_*j*_ define the peak firing rates. The excitatory influence of neuron 2 on neuron 1 is defined by

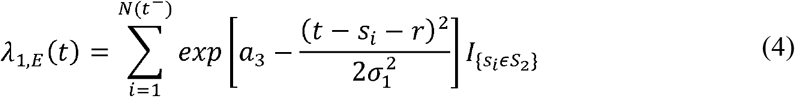

where *S*_2_ is the set containing all the spike times of neuron 2 and *r* is the time lag of the peak effect of each neuron 2 spike on neuron 1. The variable *a*_3_ defines the peak excitatory influence from neuron 2 on the firing rate of neuron 1. The history dependent terms for each neuron, are defined by

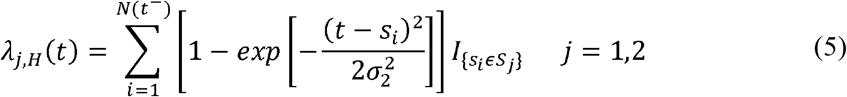

where *N*(*t*^−^) is the total number of spikes up to, but not including, time *t* and *S*_*j*_ is the set containing the spike times for each neuron. The mark process for each neuron is a scalar random variable with distribution

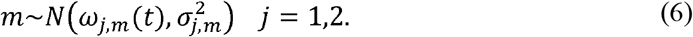

The marks are normally distributed with a time-dependent mean and a known variance 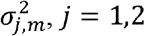. The time-dependent mean for each neuron is define by

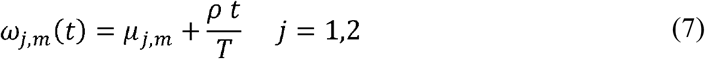

where *μ*_*j,m*_ is the time-independent component of the mean and *T* is the total time interval for the experiment. *ρ* defines how rapidly the mean of mark distribution changes as a function of time. Such a time-dependent drift in the mark could reflect changes in the spike waveform amplitude of each neuron due to electrode drift, for example. Table 3 shows the numerical values of the model free parameters. We note that these parameters are assumed to be known in both the true and mis-specified models. Here, we are focusing on assessing lack-of-fit due to model misspecification rather than due to parameter estimation error.

**Table 3:**
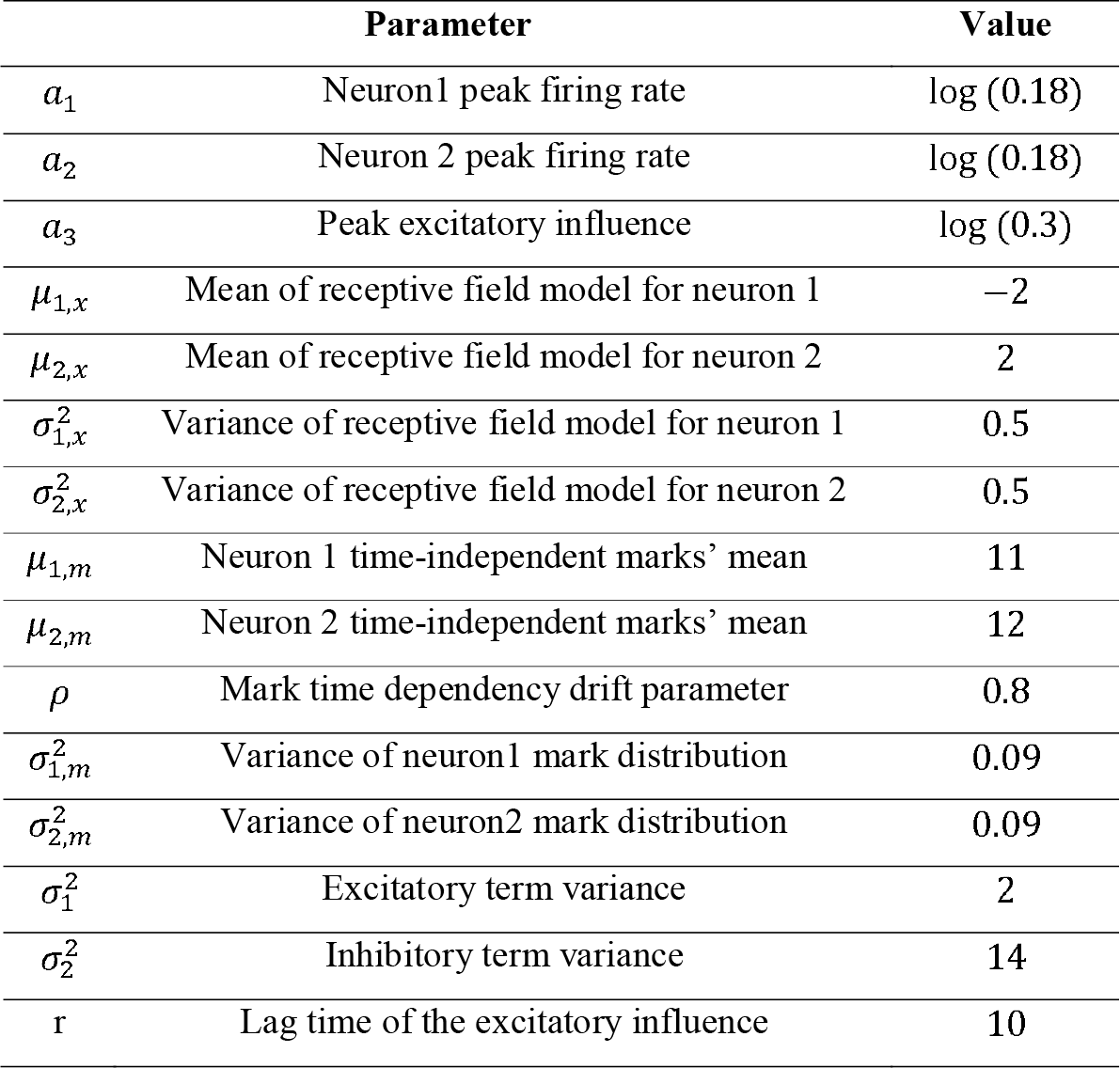
Values for the simulation model parameters

To assess how the IRCM and MDCI transformations, and the selected uniformity tests can capture the extent or lack of goodness-of-fit for marked-point process data, we generate simulated spike data using the joint marked intensity model described in Equations (2) - (7); we compare the assessed goodness-of-fit of a set of alternative models, including the true model and a number of mis-specified models, to fit this data. The true model is the one specified by Equations (2) - (7) and the parameter values in Table 3. The first mis-specified model uses the correct place and mark structure for each neuron and the interaction between them, but omits the refractory period for each neuron; the JMIF is

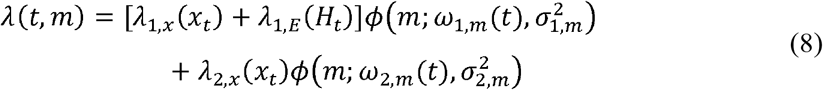

The second mis-specified model lacks only the excitatory influence of neuron 2 on neuron 1; its JMIF is

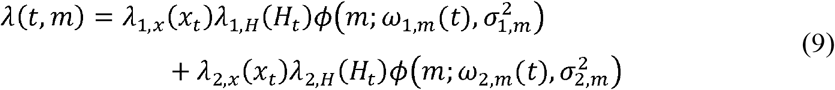

The final mis-specified model includes all components, but lacks the temporal drift in the mark distribution for both neurons; its JMIF is

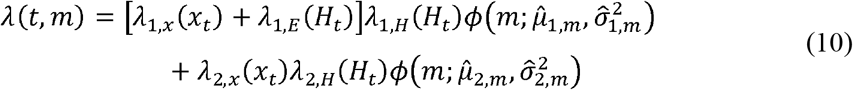

where, 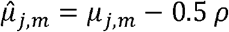 are the means and 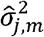 are the variances of the mark density for each neuron, based on the best estimates of these parameters using the true model under the incorrect assumption that the means are constant.

Figure 1 shows an example of the simulation data including the spike times and marks along with the position variable generated using the true model. This example includes 784 spikes - 486 from neuron 1 and 298 from neuron 2. The range of marks for neuron 1 is from 9.62 to 11.77 and for neuron 2 is from 10.53 to 12.71. This overlap means that perfect spike sorting using this single mark is impossible. Similarly, these two neurons fire over overlapping regions of space - *x*_*t*_ – as shown in Figure 1.A.

**Figure 1:**
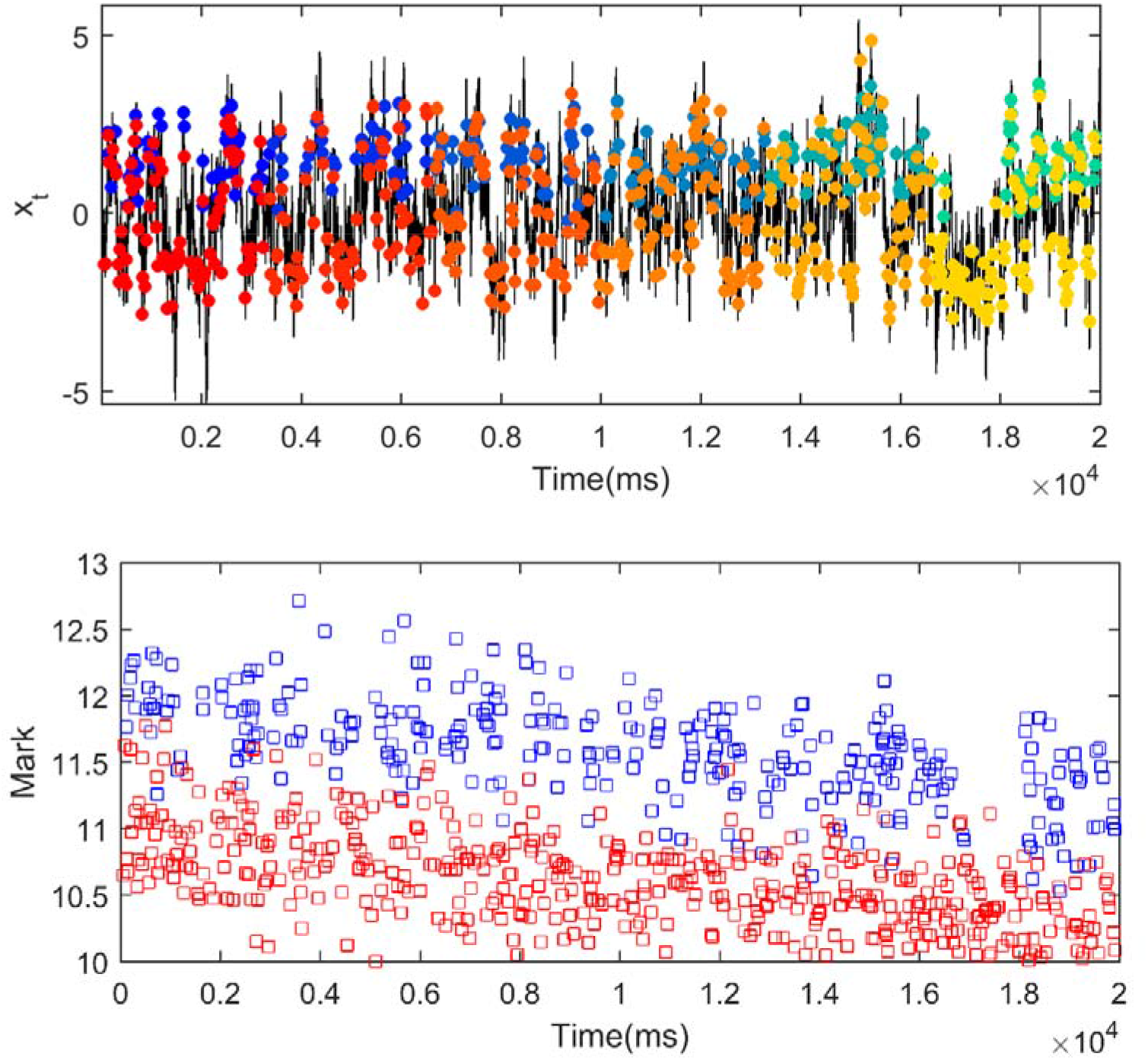
Simulated spiking by a marked-point process model with JMIF and *x*_*t*_ defined in Equations (1)-(7). There are 784 spikes in this example. **(A)** Simulated x_t_ and spike locations in time t. Spikes from neuron 1 change from red to yellow to indicate time into the simulation. Similarly, spikes from neuron 2 change from blue to green. This coloring scheme will help visualize the transformations in the IRCM and MDCI mappings in subsequent figures. **(B)** Mark values of each spike. Red and blue colors imply whether a spike comes from neuron 1 or neuron 2, respectively.

### 4.2 Transformation results

Figure 2 shows the mapping results for the IRCM transform of the simulation data using the true and alternative models. The colors of the dots indicate both the identity of the neuron generating the spike and the relative order of the spike within the experiment, matching those of Figure 1A. For the true model (Figure 2A), the transformed points are shuffled in both the rescaled time and mark axes. Visually, the transformed points appear uniformly distributed over the square; we will assess this quantitatively using multiple uniformity tests in the next section. Figure 2B shows the transformed data using the first mis-specified model, which lacks the refractory behavior of each neuron. When this inhibitory history term is omitted, the ground intensity function is over-estimated immediately after each spike, which increases the values of u. Since the missing inhibitory term does not affect the marks, the transformed data points in v axis do not show clear deviation from uniformity. Figure 2C shows the mapping result for the model missing the excitatory influence from neuron 2 to neuron 1. In this mis-specified model, a subset of the transformed data points is shifted toward lower values of u, since the intensity for neuron 1’s marks are underestimated immediately after neuron 2 spikes. In addition, since the influence of excitatory term is only on neuron 1, it is primarily the red to yellow rescaled points that are concentrated near the origin.

**Figure 2:**
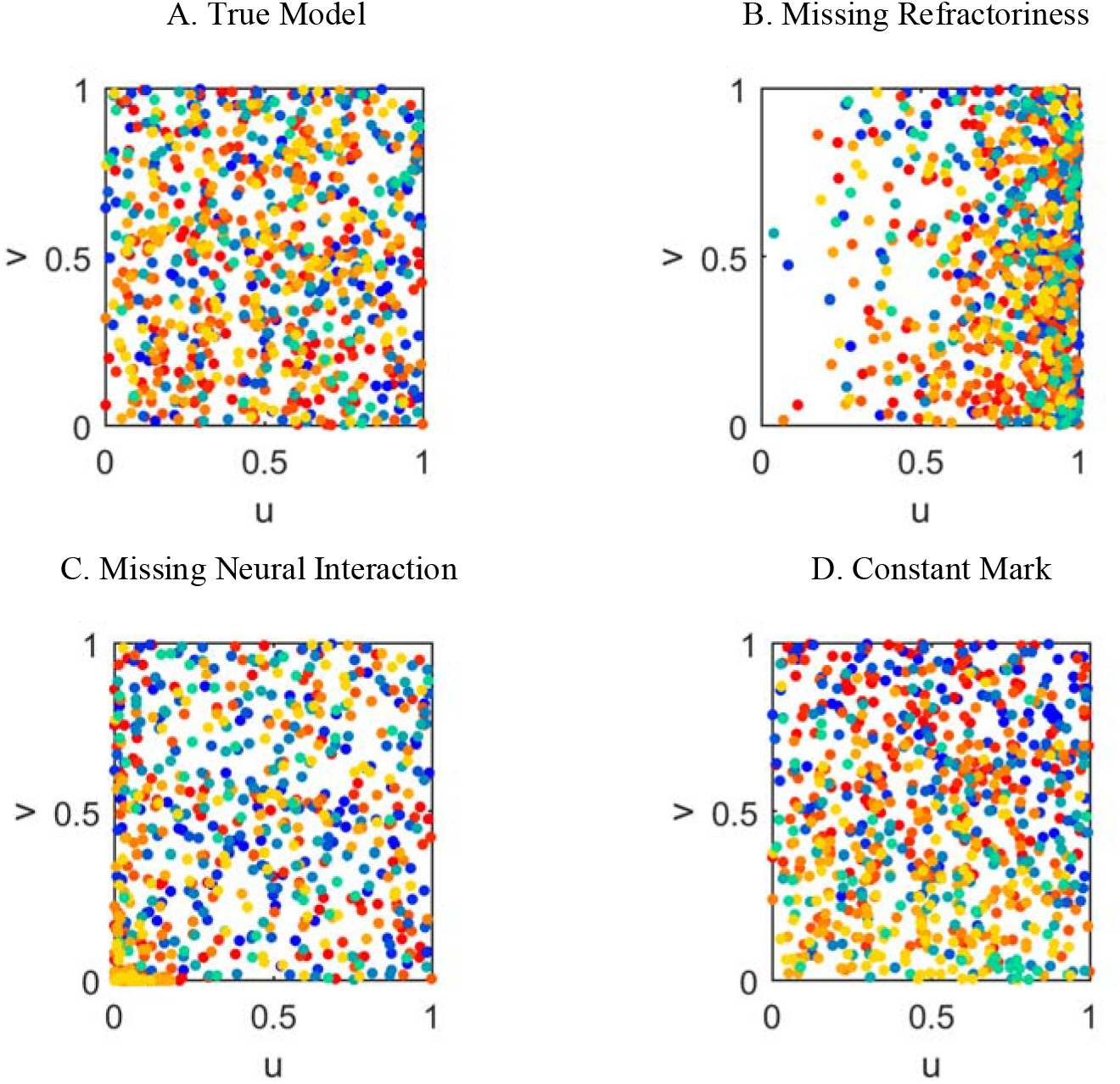
Rescaling results for the IRCM algorithm using four different models – the true model and three alternative models described in section 4.1; dot color indicates the neuron identity and timing of spike before rescaling, consistent with Figure 1A. **(A)** Rescaling using the true mark intensity function produces apparently uniform data, **(B)** Rescaling using the missing refractoriness model shows clear non-uniformity in u axis, **(C)** Rescaling using the missing interaction model shows clustering of points from neuron 1 at the origin **(D)** Rescaling using the missing mark drift model produces apparently uniform, but not independent samples

Figure 2D shows the rescaled data using the alternative model missing the drift in the mark structure. Here, there is no apparent lack of uniformity among the points, but there is a clear pattern wherein the yellow and green points from the end of the simulation session tend to cluster near the origin and the red and blue points from earlier in the session tend to cluster near the opposite corner of the square. This suggests that simple tests of uniformity might be insufficient to detect this lack-of-fit based on the IRCM transformation. In this case, including tests for independence between rescaled samples may provide a more complete view of model fit to the observed data.

Figure 3 shows the rescaling results for the MDCI algorithm. Figure 3A shows the transformed data points using the true model are distributed uniformly. In contrast to IRCM, there is little shuffling of points along the rescaled time and mark axes. In this transform, each neuron’s early spikes tend to rescale to smaller values of and its late spikes tend to rescale to later values of *u*_*i*_. Therefore, each *u*_*i*_ individually is not uniformly distributed; however, the full set of unordered rescaled times are jointly uniform.

**Figure 3:**
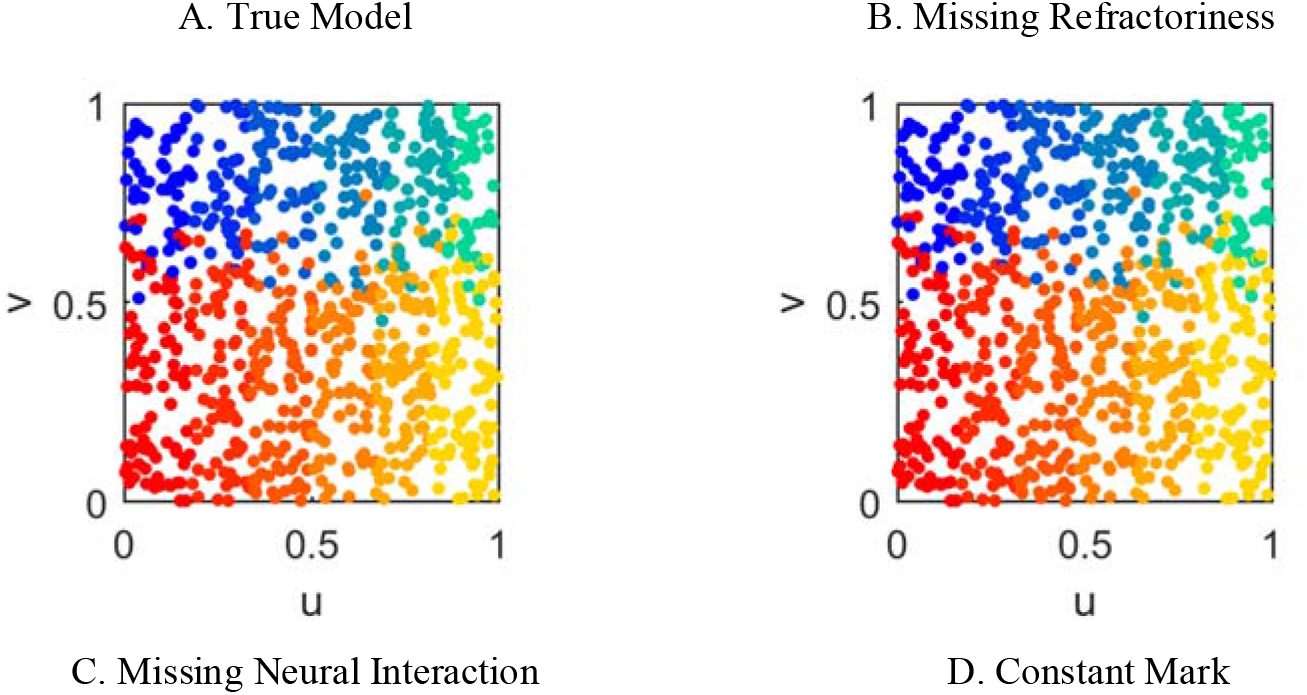
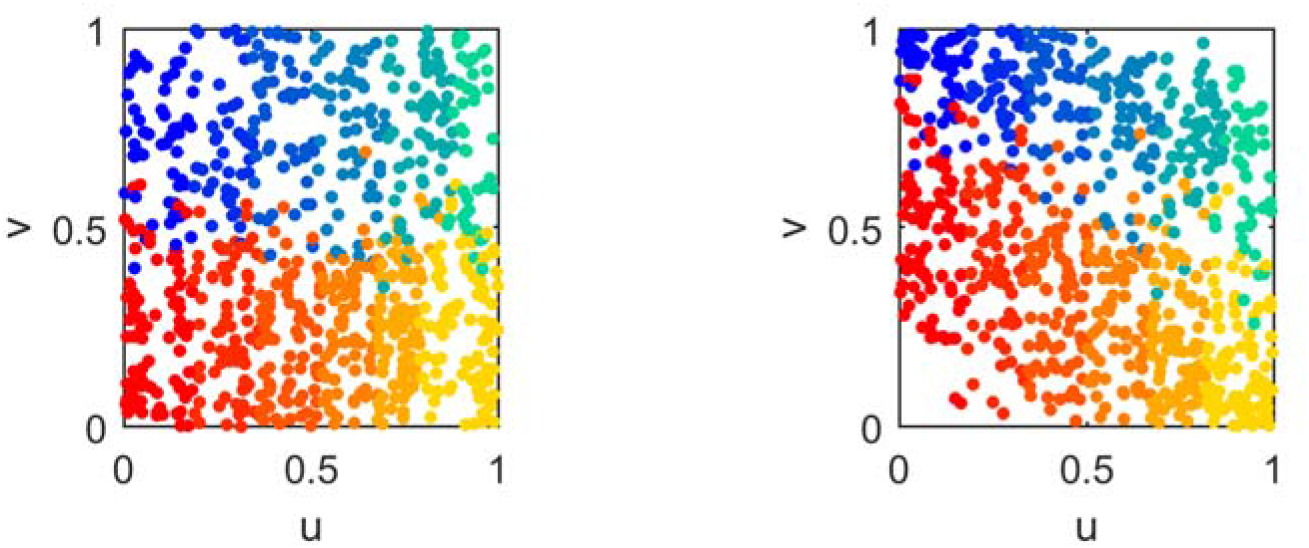
Rescaling results for the MDCI algorithm using four different models – the true model and three alternative models described in the section 4-1; dot color indicates neuron and timing of spike, consistent with Figure 1.A. **(A)** Rescaling using the true mark intensity function produces apparently uniform data, **(B)** Rescaling using missing refractoriness model appears like the true model in A. **(C)** Rescaling using the missing interaction model shows more density at lower values of v for neuron 1. **(D)** Rescaling using the missing mark drift model shows a non-uniform drift.

Figure 3B shows the rescaled points for the mis-specified model lacking refractoriness. Visually, there is no clear evidence of lack of uniformity among the samples, suggesting that tests based on this transform may lack statistical power to reject this mis-specified model. When the excitatory influence of neuron 2 on neuron 1 is omitted from the model (Figure 3C), a subtle deviation from uniformity in observed in the resulting transformed data; the fewer spikes from neuron 2 (blue to green points) occupy as much area as the more prevalent spikes from neuron 1 (red to yellow points) suggest a lack of uniformity along the axis. Figure 3D shows the rescaled points for the model lacking the drift in the marks. This leads to an apparent drift in the rescaled points, with earlier spikes producing larger values of v and later spikes producing smaller values of v. Unlike the IRCM transformation, the lack of uniformity is visually clear for this mis-specified model.

Figures 2 and 3 suggest that different forms of model mis-specification may be better identified using different transformations; the missing refractoriness model shows clear lack-of-fit based on the IRCM but not the MDCI transformation, while the missing mark drift model shows more apparent lack-of-fit through the MDCI transformation. It remains to be seen whether this apparent lack-of-fit is captured quantitatively using each of the uniformity tests described previously; we explore this in the following Section.

### 4.3 Uniformity Test Results

Tables 4 and 5 provide the results of the uniformity tests described in Table 2 along with their corresponding -values on the rescaled data shown in Figures 2 and 3 using the IRCM and MDCI transformations. A small p-value indicates strong evidence against the null hypothesis; here, we set the significance level *α* of 0.05. The null hypothesis is that the sample data are distributed uniformly in a unit square; this hypothesis would be true if the original marked point process data are generated based on the joint mark intensity model used for the transformation.

**Table 4:**
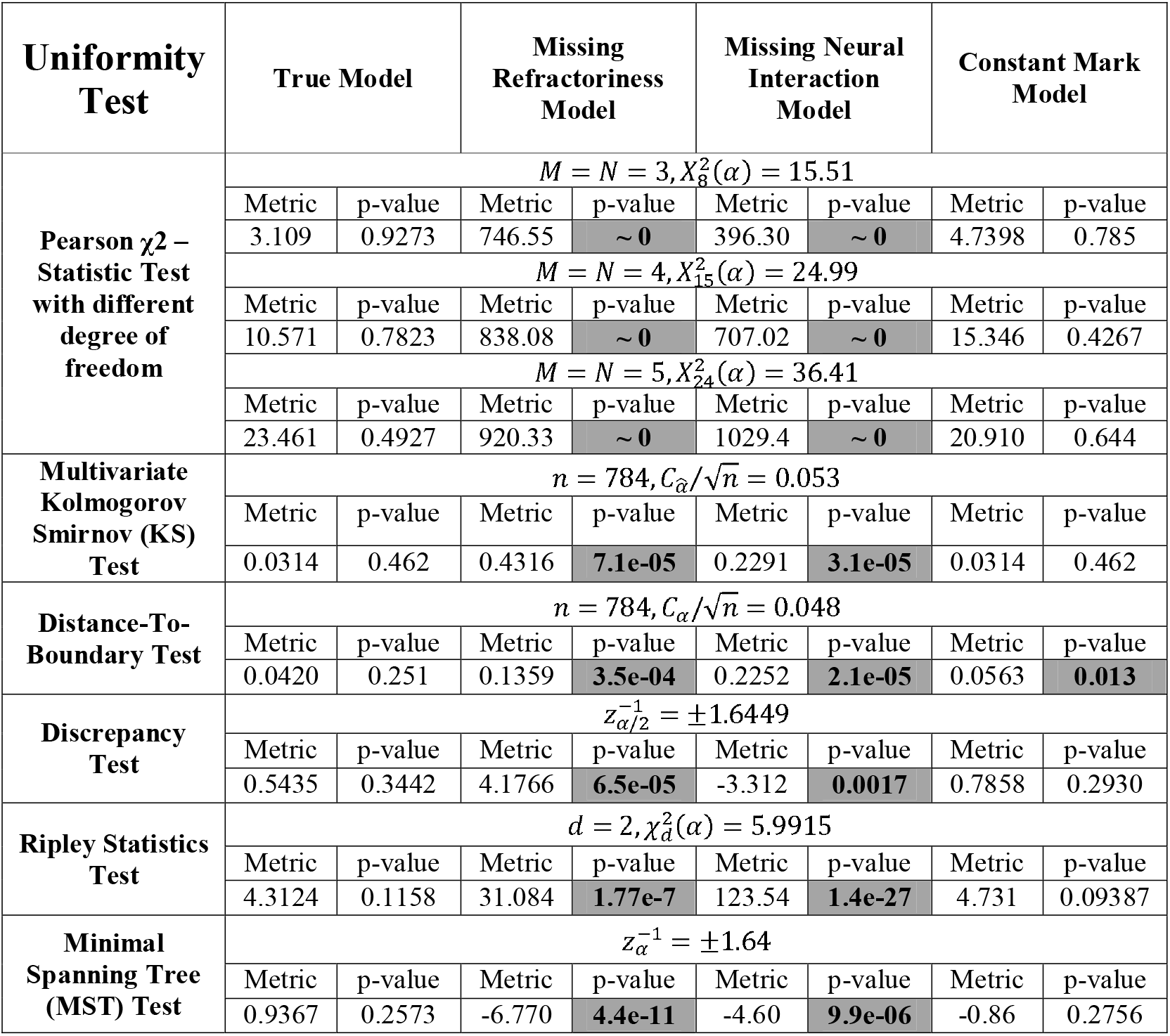
Different uniformity test statistics and corresponding *p*-values using the IRCM algorithm; the bold numbers (gray boxes) show cases where the test identified lack of fit at a significance-level of *α* = 0.05

**Table 5:** Different uniformity test statistics and corresponding p-values using the MDCI algorithm; the bold numbers (gray boxes) show cases where the test identified lack of fit at a significance-level of *α* = 0.05

| Uniformity<br>Test | True Model |  | Missing<br>Refractoriness<br>Model |  | Missing Neural<br>Interaction<br>Model |  | Constant Mark<br>Model |  |
| --- | --- | --- | --- | --- | --- | --- | --- | --- |
| Pearson $\chi^2$ –<br>Statistic Test<br>with different<br>degree of<br>freedom | $M = N = 3, X_8^2(\alpha) = 15.51$ | | | | | | | |
|  | Metric | p-value | Metric | p-value | Metric | p-value | Metric | p-value |
|  | 1.997 | 0.981 | 2.788 | 0.946 | 54.33 | 5.9e-9 | 94.510 | ~ 0 |
| | $M = N = 4, X_{15}^2(\alpha) = 24.99$ | | | | | | | |
|  | 10.22 | 0.805 | 10.693 | 0.774 | 77.87 | 1.7e-10 | 117.79 | ~ 0 |
| | $M = N = 5, X_{24}^2(\alpha) = 36.41$ | | | | | | | |
|  | Metric | p-value | Metric | p-value | Metric | p-value | Metric | p-value |
|  | 21.202 | 0.626 | 18.423 | 0.782 | 82.900 | 2.08e-8 | 139.15 | ~ 0 |
| Multivariate | $n = 784, C_{\hat{a}}/\sqrt{n} = 0.053$ | | | | | | | |
| **Kolmogorov Smirnov Test** | Metric | p-value | Metric | p-value | Metric | p-value | Metric | p-value |
|  | 0.038 | 0.307 | 0.0152 | 0.834 | 0.120 | **9.1e-04** | 0.079 | **0.007** |
| **Distance-To-Boundary Test** | $n = 784, C_\alpha/\sqrt{n} = 0.048$ | | | | | | | |
|  | Metric | p-value | Metric | p-value | Metric | p-value | Metric | p-value |
|  | 0.0162 | 0.814 | 0.023 | 0.661 | 0.035 | 0.368 | 0.028 | 0.524 |
| **Discrepancy Test** | $z_{\alpha/2}^{-1} = \pm 1.6449$ | | | | | | | |
|  | Metric | p-value | Metric | p-value | Metric | p-value | Metric | p-value |
|  | -0.209 | 0.390 | -0.169 | 0.393 | 0.245 | 0.387 | 0.037 | 0.398 |
| **Ripley Statistics Test** | $d = 2, \chi_d^2(\alpha) = 5.9915$ | | | | | | | |
|  | Metric | p-value | Metric | p-value | Metric | p-value | Metric | p-value |
|  | 1.862 | 0.394 | 2.376 | 0.304 | 3.920 | 0.140 | 4.903 | 0.086 |
| **Minimal Spanning Tree (MST) Test** | $z_\alpha^{-1} = \pm 1.64$ | | | | | | | |
|  | Metric | p-value | Metric | p-value | Metric | p-value | Metric | p-value |
|  | 0.128 | 0.395 | -0.150 | 0.394 | -1.341 | 0.162 | -2.561 | **0.0150** |

The results presented in Tables 4 and 5 suggest that there is no single combination of transform and uniformity test that will identify all forms of model lack-of-fit. For this simulation, the IRCM transformation makes it simple to identify lack of fit due to incorrect history dependence structure – either missing refractoriness or neural interactions – using any of the uniformity tests. However, it remains difficult to detect lack-of-fit due to missing the mark dynamics; while the distance-to-boundary test detects the mis-fit at the *α* = 0.05 significance level, this result would not hold up to correction for multiple tests. The MDCI transformation is not able to detect mis-fit in the missing refractoriness model using any of the uniformity tests, but both the Pearson and multivariate KS tests are able to detect lack of fit due to missing neural interactions and missing mark dynamics at very small significance levels.

These results also suggest that certain uniformity tests may achieve substantially higher statistical power over others for the types of lack-of-fit often encountered in neural models. While all of the tests were able to identify the mis-specified models missing history dependent components via the IRCM transformation, the Pearson and multivariate KS tests provided much lower p-values for detecting the missing interaction and constant mark models’ mis-fit under the MDCI transformation. This suggests that different combinations of transformations and uniformity and independence tests can provide different views on goodness-of-fit that can be used together to provide a more complete picture of the quality of a model.

While the IRCM transformed samples in the constant mark model do not show obvious lack of uniformity in Fig. 2D, these samples do show obvious dependence – as seen through the structure in the dot colors. The rescaling theorem for this transformation guarantees that under the true model, these samples will be independent. We can therefore apply correlation tests to these samples to further assess model goodness-of-fit. To demonstrate, we used a correlation test between consecutive samples based on a Fisher transform [30], defined by

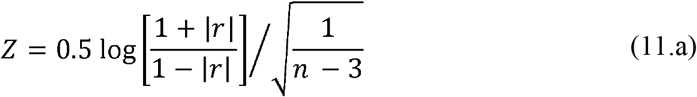

where *r* is the correlation coefficient between *v*_*i*_ and *v*_*i*+1_ and *n* is the number of samples. Under the true model, the p-value for this test is 0.58 suggesting no lack-of-fit related to dependence in the transformed samples; under the constant mark model, the p-value was 2.1e-08, suggesting lack of fit in the model leading to dependence of the samples. This suggests that uniformity and independence tests can provide complementary tools to identify model misspecification in IRCM transformed samples.

## 5 Discussion

A fundamental component of any statistical model is the ability to evaluate the quality of its fit to observed data. While the marked point process framework has the potential to provide holistic models of the coding properties of a neural population while avoiding a computationally expensive spike-sorting procedure, until recently methods for assessing their goodness-of-fit have been lacking. Our preceding work to extend the time-rescaling theorem [5] to marked point process neural models has provided a preliminary approach to address this problem [6] but further work was necessary to make the approach computationally efficient in higher dimensions, to enable the use of more statistically powerful test methods, and to understand which tests are most useful for capturing different aspects of model misspecification.

In this paper, we proposed two new transformations – IRCM and MDCI – that combine rescaling in both the time and mark spaces, to produce samples that are uniformly distributed in a hypercube for the correct marked point process model. This removes one of the most troublesome issues with our prior method, the fact that time-rescaling produced uniform samples in a random space that could be computationally challenging to compute, precluded multiple uniformity tests, and made those tests that could be performed more computationally challenging. In particular, these methods can reduce concerns in designing population coding models that using high dimensional spike features will make model assessment intractable; instead the focus of waveform feature selection for these models can be on finding the features that best explain the population coding properties.

While both the IRCM and MDCI transformations produce samples that are uniformly distributed in a hypercube for the true model, each transformation can capture different attributes of the quality of the model fit to the observed data. The IRCM rescales the inter-spike intervals between all observed spikes, irrespective of their waveforms, and then rescales the marks in a time-dependent manner. For correct models, this causes mixing between the spike marks from different neurons. This transformation is likely to be particularly sensitive to misspecification of interactions between neurons as in our simulation example. The MDCI transformation rescales the spike waveform features irrespective of when they occur and then rescales the spike times in a manner that depends upon their waveforms. This transformation tends to keep spikes from a single neuron nearby, and is likely to be sensitive to misspecification of the coding properties of individual neurons. The fact that the IRCM makes the rescaled samples independent allows us to use correlation tests as further goodness-of-fit measures. The fact that the MDCI keeps marks from individual neurons nearby allows us to identify regions of nonuniformity in the hypercube to determine which waveforms have spiking that is poorly fit by the model. Together, these transformations provide complimentary approaches for model assessment and refinement.

In addition to having multiple, complimentary transformations for the data, we have multiple tests for uniformity and dependence with which to assess the transformed samples. Here, we explored six well-established uniformity tests to examine how different forms of model misspecification could be captured using combinations of these transforms and tests. As expected, the true model did not lead to significant lack of uniformity in either transformation based on any of the tests we explored. Similarly, for the true model, our correlation test did not detect dependence in the IRCM transformed samples. For the misspecified models, different combinations of transformations and uniformity tests were able to identify different sources of lack-of-fit. The missing refractoriness and missing neural interaction models were easily identified as mis-fit under the IRCM transform using all of our tests, but the constant mark model could not be identified by any of the tests using this transform. The constant mark model was identified as mis-fit under the MDCI using the Pearson and multivariate KS tests but not the other uniformity tests. Across these simulations, the Pearson Chi-Square, Multivariate KS, and MST tests proved to be statistically more powerful in capturing the particular forms of model misspecification that we examined. However, these simulations were limited both by using a simple two-neuron population model and by using only a one dimensional mark. While more systematic exploration of uniformity tests are necessary to know which combinations of transforms and tests are most useful for determining different aspects of model goodness-of-fit, these results suggest that no one combination is likely to work in all cases. Relatedly, goodness-of-fit for marked point process models should not be limited to rescaling methods; deviance analysis and point process residuals can provide additional, complementary goodness-of-fit measures. A toolbox that includes multiple approaches, including different rescaling transformations and tests provides substantially more statistical power than any one approach on its own.

Ultimately, insight into which goodness-of-fit methods are most useful for these clusterless coding models will require extensive analysis of real neural population spiking data. Based on the many advantages of the clusterless modeling approach – the reduction of bias in receptive field estimation [31], the ability to use spikes cannot be sorted with confidence [2], the ability to fit models in real time for during the recording sessions – and the experimental trend toward recording larger populations and closed-loop experiments, we anticipate that clusterless modeling approaches and methods to assess their quality will become increasingly important. In order to enable experimentalists to apply these algorithms in their data analysis, we have made the MATLAB code for these transformations along with the uniformity tests explored here available through our Github repository at https://github.com/YousefiLab/Marked-PointProcess-Goodness-of-Fit

## Appendix A. Rescaling Theorem Proofs

In this paper, we introduced IRCM and MDCI algorithms. In this section we present the theoretical proof for these algorithms in two separate subsections.

## Appendix A.1. Interval-Rescaling Conditional Mark Distribution (IRCM)

In this section, we provide a set of new transformations and their properties in transforming observed marked-point process data points to uniformly distributed samples in hypercube. We take different methodologies to prove properties of these transformations; we either use change of variables’ theorem [32] or derive the distribution of observed joint mark and spike events under these transformations in these proofs. We assume that we have a sequence of marked-point process with observed marks 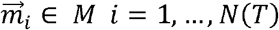 associated with the spike time 0 = *S*_0_ < *S*_1_ < *S*_2_ < ··· < *S*_*i*_ < ··· < *S*_*N*(*T*)_ < *T* and with a joint mark intensity function 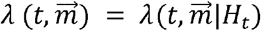. The joint probability of observing *N*(*T*) events over the period of *t* = [0 *T*] is defined by

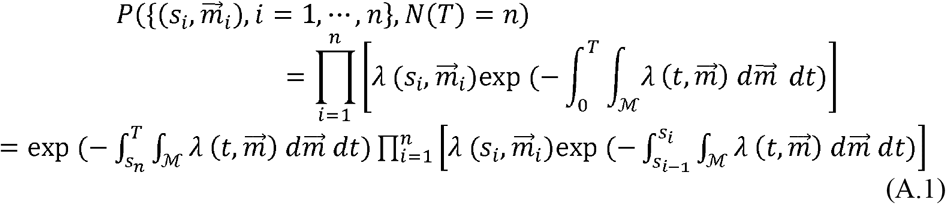

and we show the following transformation takes 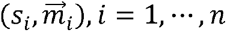 to a set of new data points (*u*_*i*_, *v*_*i*_), *i* = 1, ···, *n*. which are i.i.d samples with a uniform distribution in the range of [0 1]^1+*d*^ − *d* is the dimension of mark.

### Theorem 1

Let’s define the ground conditional intensity by

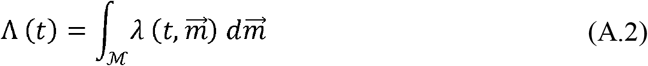

and conditional intensity of mark given the event time by

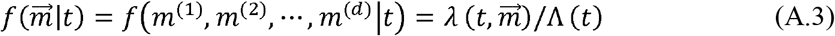

where m^(l)^ is *l*^*th*^ element of the vector 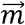. The conditional intensity of mark can be written by

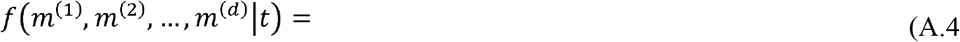

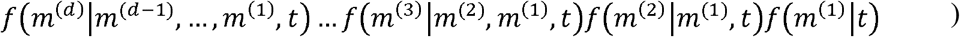

where,

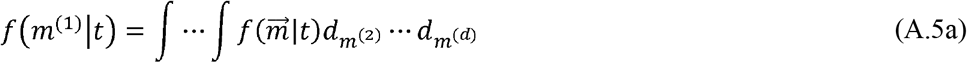

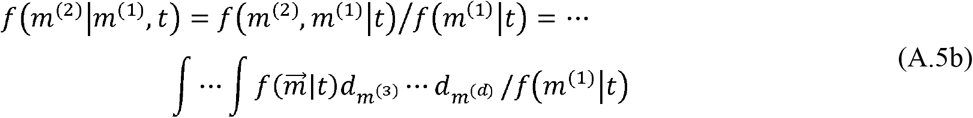

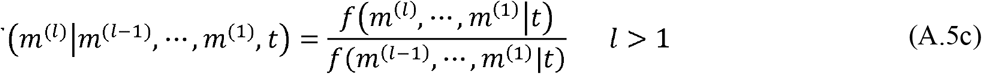

Now, we can define the following *d* + 1 new variables 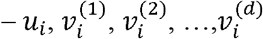, 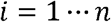:

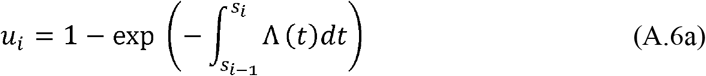

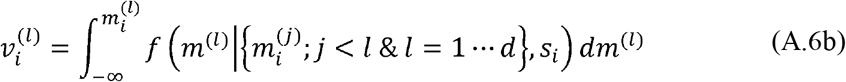

which are i.i.d. samples with a uniform distribution in [0 1]^*d*+1^ hypercube, under the true model.

**Proof:** We can redefine Equation (A.1) by,

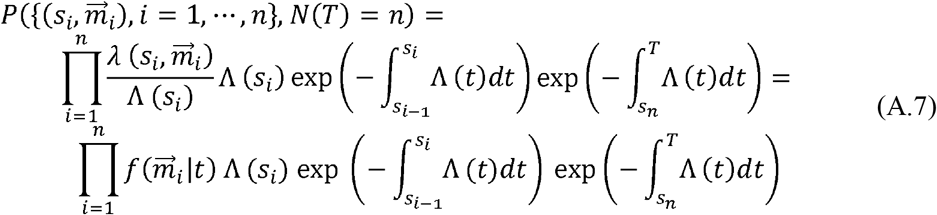

here, the first n terms represent the probability of observing continuous samples over the time period of [0, *s*_*n*_] with marks 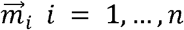. The last term corresponds to the probability of not observing any event for the time period of (*s*_*n*_; *T*]; this corresponds to *N*(*T*) − *N*(*s*_*n*_) = 0.

We want to build the joint probability distribution of 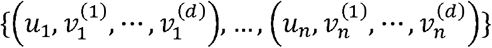 given 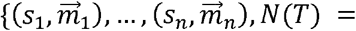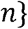. We first focus on the time period [0, *s*_*n*_], where we observe n events. We use the change of variable theorem [32] to build the joint probability distribution of full events over this time period. To derive the joint probability distribution, we need to calculate the Jacobian matrix, which is defined by

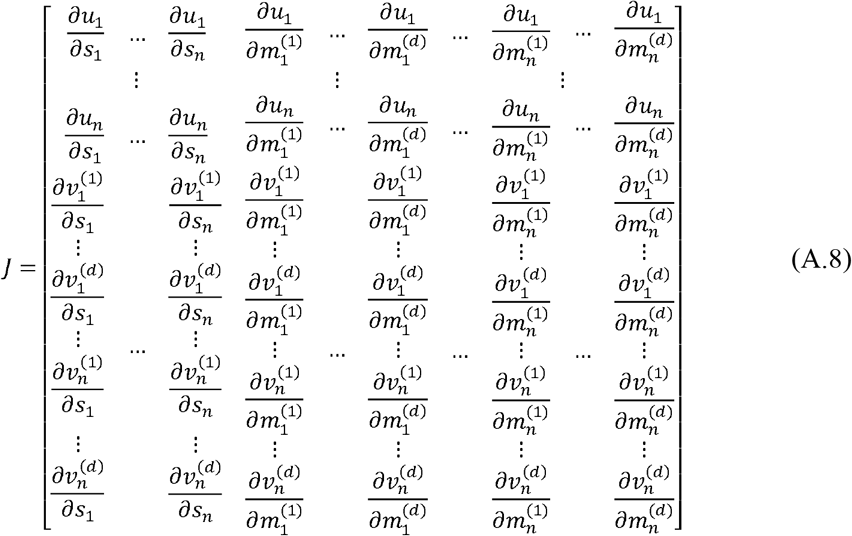

where,

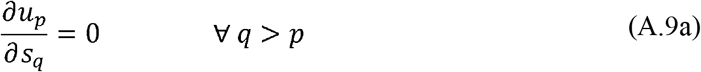

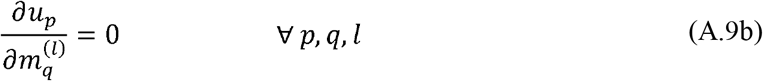

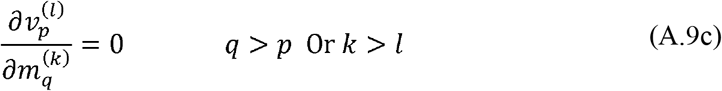

where, *p*, *q* ∈ {1,…,*n*} and *l*, *k* ∈ {1,…,*d*}. Note that, the upper triangular elements of the matrix J are equal to zero, and thus we only need to calculate the diagonal elements of the matrix to calculate its determinant. The matrix diagonal elements are defined by

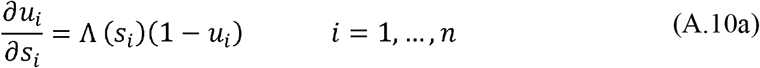

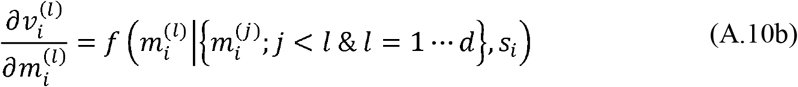

The Jacobian matrix determinant is equal to

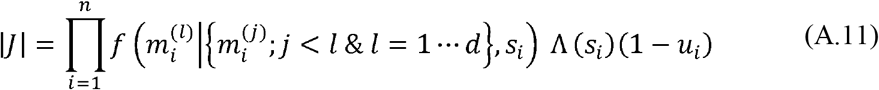

by replacing these elements in the Equation (A.7), we get

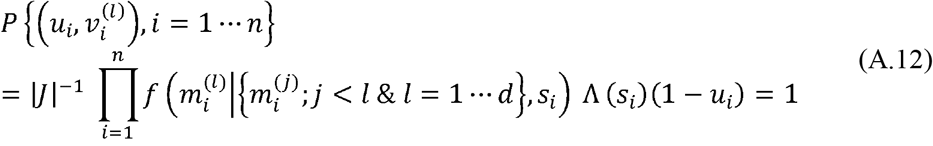

Now, we consider the last component of the distribution which implies no event from time s_n_ to T. Let’s define a new random variable z,

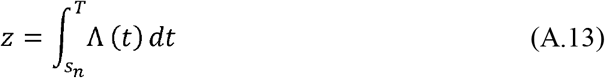

which defines the number of events for the time s_n_ to T. We define the probability of not observing an event by

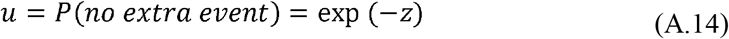

where, u is a new random variable in the range of 0 to 1. By changing the variable from z to u, the joint probability distribution can be written by

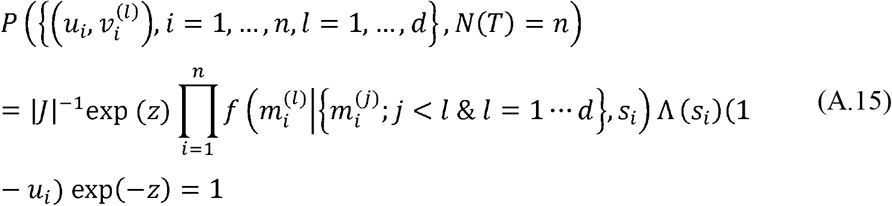

Thus, all elements including the last element are uniformly distributed on [0 1]^*d*+1^ given the assumption that samples are generated using the true 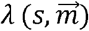.

Not that the last element becomes a sure event in u space. The last term can be also projected back to u_i_s; with this assumption – when it is not compensated – the transformed u_i_s are uniformly distributed on a scaled space of 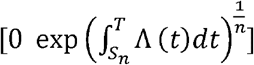.

### Corollary 1

The marginalization steps over the mark dimension described in Theorem 1 is valid on any arbitrary sequence of the mark space dimension.

### Corollary 2

The u_i_ samples generated by Equation (A.6a) given the ground conditional intensity defined in Equation (A.2) are independent and uniformly distributed over the range 0 to 1 independent of v samples. samples are independent of v samples, and the mapping over u corresponds to a time rescaling theorem [5] over the full event’s time intervals.

## Appendix A.2. Mark Density Conditional Intensity (MDCI)

In this section, we provide a proof for Algorithm 2 using the following theorem.

### Theorem 2

Let’s assume the following pdfs are defined using *λ*(*t*, *m*),

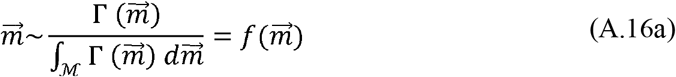

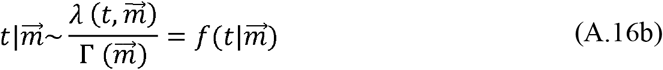

where, 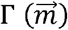, is defined by

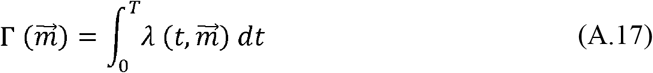

Now, we can define the following *d* + 1 new variables 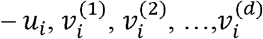, 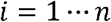,

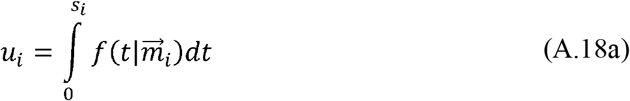

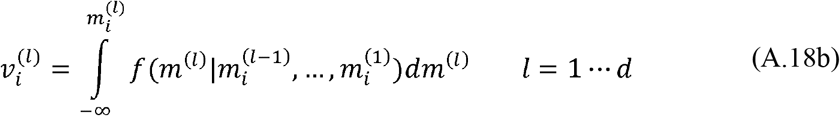

which are rescaled samples with a uniform distribution in [0 1]^*d*+1^ hypercube, under the true model.

**Proof:**

The joint probability distribution of full event defined in equation (A.1) can be rewritten by

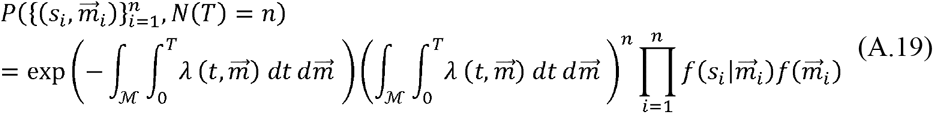

To prove the theorem, we require to build the joint probability distribution of 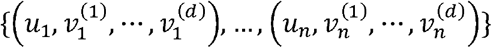 given 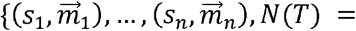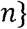. First, we define 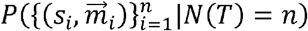, which corresponds to

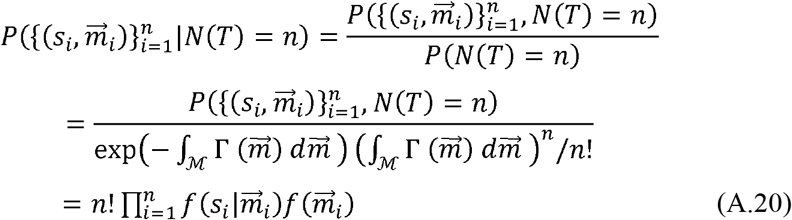

In equation (A.20), the denominator defines the joint probability distribution of observing *n* events independent of their temporal order. Given the history dependence of the events, the joint probability distribution of temporally ordered events [6] is defined by

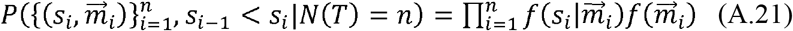

In equation (A.21), 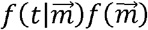 defines the joint probability distribution of the mark and spike time. Equation set (A.18) is the Rosenblatt Transformation [7] of the spike time and mark, mapping the observed events from multivariate continuous random variables defined by 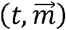 to another one, 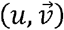. Under the Rosenblatt Transformation theorem, the transformed data points are uniformly distributed in a hypercube of [0 1]^*d*+1^. As a result, the joint distribution of 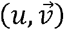 is define by

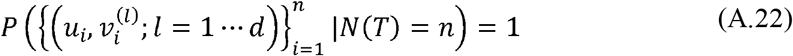

which suggests that 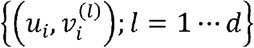 s are uniform samples in a hypercube of [0 1]^*d*+1^

## Appendix B. 1. Mark-Rescaling Conditional Intensity (MRCI)

The MRCI algorithm starts by building the mark intensity function – Γ(*m*), which is followed by deriving the conditional intensity function – *f*(*t*|*m*_*i*_). This transformation corresponds to a time-rescaling on the mark and the cdf of conditional intensity on spike time. The table below describes the steps being taken to map the full spike event data to a unit square.

### Algorithm 3

Mark-Rescaling Conditional Intensity (MRCI)

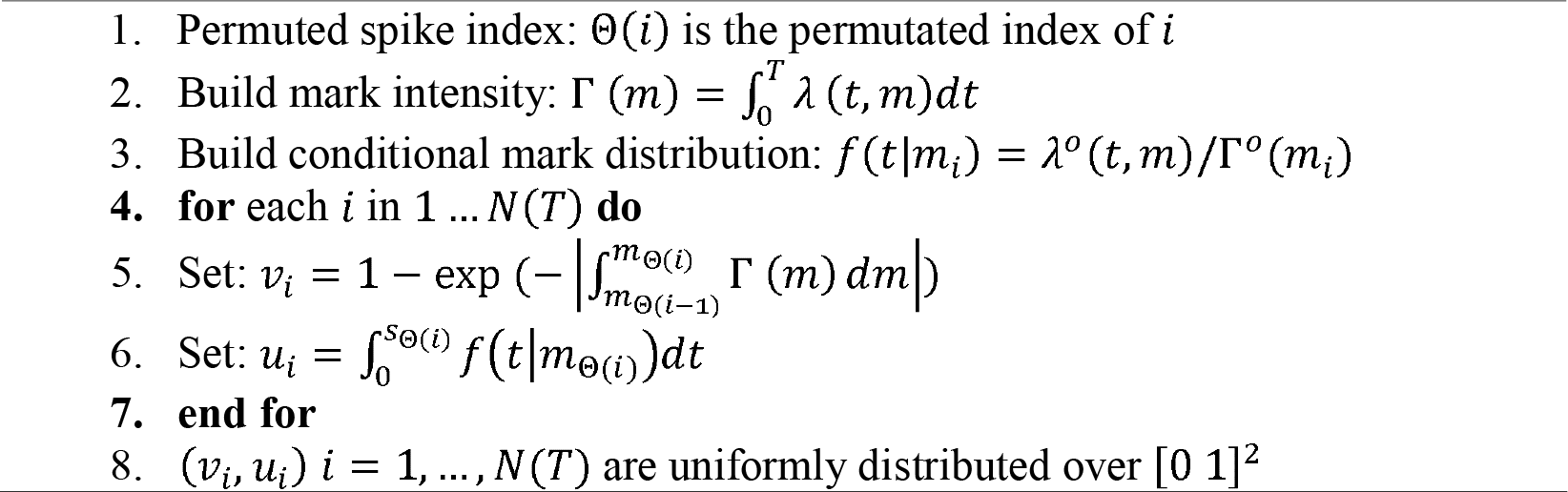

In MRCI algorithm, we can use any permutated sequence over mark and the resulting (*v*_*i*_, *u*_*i*_) samples still hold uniformity. We provide the theoretical proof of MRCI transformation in Appendix B.2 section. Like the previous algorithms, we can use multiple uniformity tests, described in section 3, to assess the accuracy of model fit.

In contrast to IRCM and MDCI, where we marginalized the conditional mark distribution over the mark space, marginalizing of Γ(*m*) to construct uniformly distributed variables is not easy. When the mark is a scalar variable, we can use u_i_ and v_i_ samples to draw further insight about the model fit. u_i_ samples, when they are constructed using sorted marks, reflect how properly the temporal properties of the full events are captured using the model.

A challenge with MRCI, when the observed mark is multidimensional, is the interpretation of full spike events in the multi-dimensional spaces. There is no unique solution on how we should sort multi-dimensional marks. As a result, MRCI can be a proper transformation under two circumstances: a) when the mark, independent of its dimension, is treated as one element, b) when the dimension of the mark is one.

We utilize the simulation data described in section (4-1) to assess the mapping result of MRCI. Figure A1 shows the mapping results for this algorithm. For the true model, we expect the observed data points are being mapped to uniformly distributed samples in a unit square - Figure A1. A. The results are in a strong support of this assumption, and Figures A1. B to A1. D show the mapping results using alternative models under MRCI mapping. For the inhibitory independent model, the transformed data points are shifted toward higher values in axis. This is because, when the history term is dropped, the joint mark intensity function is over-estimated for the times after each spike and thus the mark intensity function increases in the interval between pair of events - check lines 5 in MRCI algorithm. Similarly, for excitatory independent model, the new data points are shifted toward low values of axis; this is because by eliminating excitatory effect of neuron 2 on neuron 1, the joint mark intensity function is getting under-estimated for the time after neuron 2 spike and thus the mark intensity function decreases in the interval between pair of events - check lines 5 in MRCI algorithm. Figure A1. D shows the transformed data points under time independent mark model. Like the IRCM, the MRCI algorithm is not capable of capturing the misspecification being embedded in the mark distribution; this is because we take integral of the mark intensity function in interval between pair of events where the mark intensity function simultaneously increases and decreases over theses intervals.

**Figure A1:**
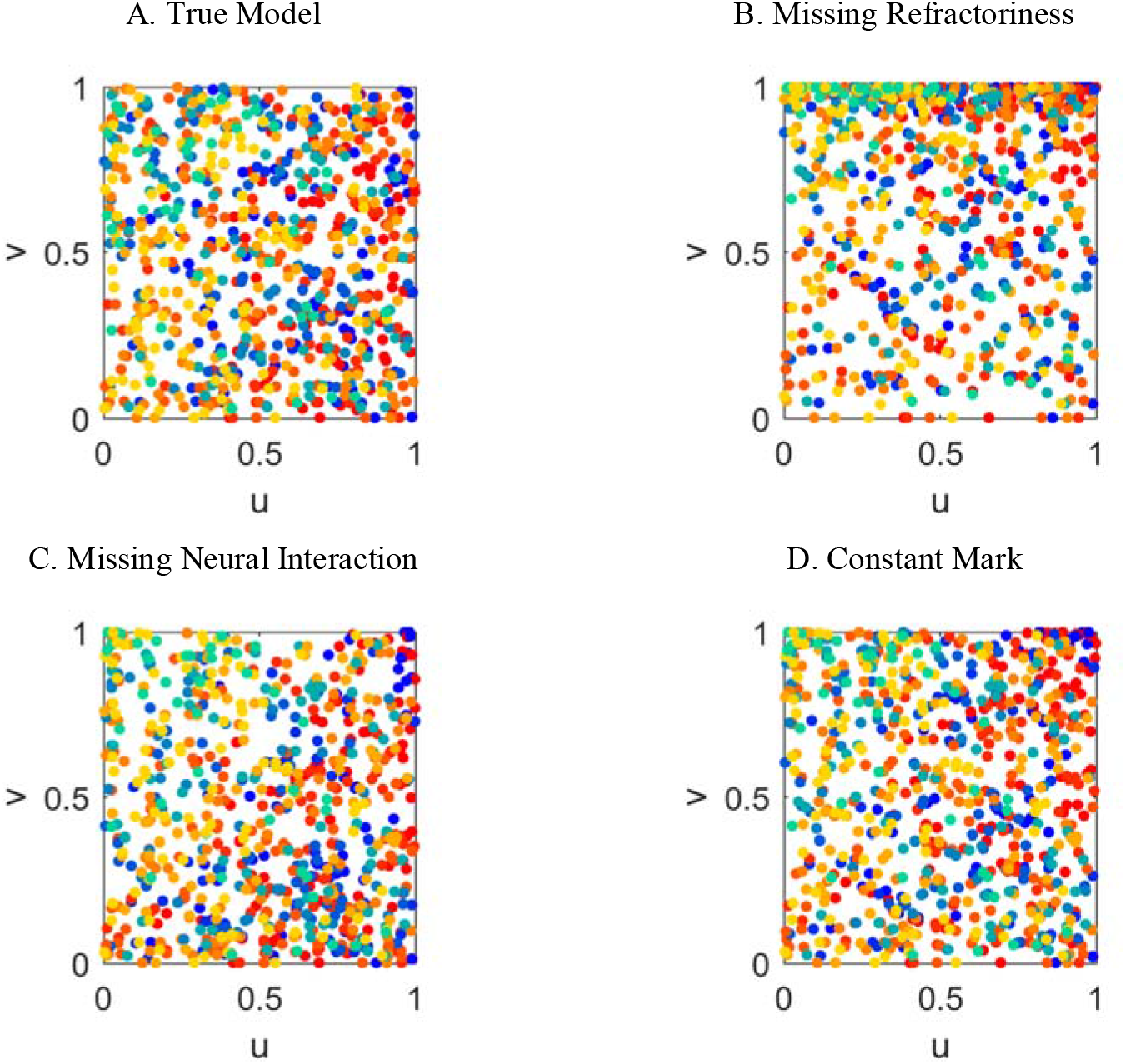
Rescaling results for the MRCI algorithm using four different models – the true model and three alternative models described in the section 4-1; dot color indicates neuron and timing of spike, consistent with Figure 1.A. **(A)** Rescaling using the true mark intensity function produces apparently uniform data, **(B)** Rescaling using missing refractoriness model shows a non-uniform drift. **(C)** Rescaling using the missing interaction model shows more v. **(D)** Rescaling using the missing mark drift model appears like the true model in A.

The uniformity test results for the MRCI algorithm are reported in the Table A1. Given the result in the table, Chi-Square Pearson and Multivariate KS tests reject the null hypothesis for all alternative models. The Distance-To Boundary, Discrepancy, Ripley Statistics and MST tests only reject the null hypothesis for inhibitory independent model and fail to reject the null hypothesis for other miss-specified models. The results here are in accordance to the previous result in Tables 5 and 6, where Chi-Square Pearson and Multivariate KS tests show to be statistically stronger tests. The statement that we should try a combination of uniformity tests to build a stronger confidence in the goodness-of-fit result holds for this transformation as well.

**Table 1.A:** Different uniformity tests’ metrics and corresponding p-values applied on the data points transformed using MRCI algorithm; the bold numbers (gray boxes) show cases that the null hypothesis is rejected with a significance-level of *α* = 0.05

| Uniformity Test | True Model |  | Missing Refractoriness Model |  | Missing Neural Interaction Model |  | Constant Mark Model |  |
| --- | --- | --- | --- | --- | --- | --- | --- | --- |
| Pearson $\chi^2$ – Statistic Test with different degree of freedom | $M = N = 3, X_8^2(\alpha) = 15.51$ | | | | | | | |
|  | Metric | p-value | Metric | p-value | Metric | p-value | Metric | p-value |
|  | 8.6071 | 0.3765 | 279.07 | ~0 | 44.452 | 4.65e-7 | 33.484 | 5.03e-5 |
| | $M = N = 4, X_{15}^2(\alpha) = 24.99$ | | | | | | | |
|  | Metric | p-value | Metric | p-value | Metric | p-value | Metric | p-value |
|  | 15.931 | 0.3867 | 357.77 | ~0 | 53.061 | 3.7e-6 | 45.469 | 6.45e-5 |
| | $M = N = 5, X_{24}^2(\alpha) = 36.41$ | | | | | | | |
|  | Metric | p-value | Metric | p-value | Metric | p-value | Metric | p-value |
|  | 22.452 | 0.5523 | 453.36 | ~0 | 67.658 | 4.92e-6 | 74.035 | 5.26e-7 |
| Multivariate Kolmogorov Smirnov Test | $n = 784, C_{\bar{\alpha}}/\sqrt{n} = 0.053$ | | | | | | | |
|  | Metric | p-value | Metric | p-value | Metric | p-value | Metric | p-value |
|  | 0.0510 | 0.0510 | 0.2974 | 7.25e-5 | 0.0857 | 0.003 | 0.0820 | 0.005 |
| Distance-To Boundary Test | $n = 784, C_{\alpha}/\sqrt{n} = 0.048$ | | | | | | | |
|  | Metric | p-value | Metric | p-value | Metric | p-value | Metric | p-value |
|  | 0.0282 | 0.536 | 0.1843 | 2.29e-4 | 0.035 | 0.366 | 0.0432 | 0.232 |
| Discrepancy Test | $z_{\alpha/2}^{-1} = \pm 1.6449$ | | | | | | | |
|  | Metric | p-value | Metric | p-value | Metric | p-value | Metric | p-value |
|  | -0.667 | 0.3193 | -1.720 | 0.0408 | -0.893 | 0.2677 | -1.351 | 0.1602 |
| Ripley Statistics Test | $d = 2, \chi_d^2(\alpha) = 5.9915$ | | | | | | | |
|  | Metric | p-value | Metric | p-value | Metric | p-value | Metric | p-value |
|  | 2.3306 | 0.3118 | 23.374 | 8.4 e-9 | 4.2077 | 0.122 | 4.5260 | 0.104 |

| Minimal<br>Spanning Tree<br>(MST) Test | $z_{\alpha}^{-1} = \pm 1.6449$ | | | | | | | |
| --- | --- | --- | --- | --- | --- | --- | --- | --- |
|  | Metric | p-value | Metric | p-value | Metric | p-value | Metric | p-value |
|  | 0.9616 | 0.2513 | -5.133 | **7.5e-07** | -1.492 | 0.1311 | -1.111 | 0.2151 |

## Appendix B.2. Mark-Rescaling Conditional Intensity (MRCI) Theoretical Proof

### Theorem 3

Let’s define the ground conditional intensity over the mark space by:

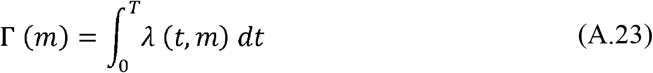

and conditional intensity of the event given the mark by

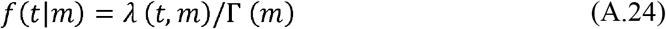

Then, the following two new variables – u_i_ and v_i_:

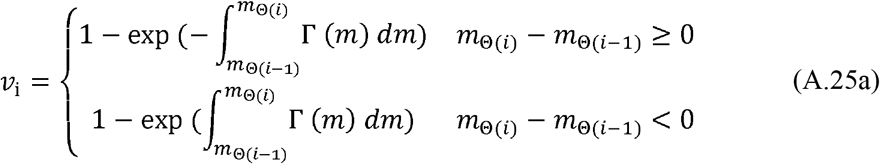

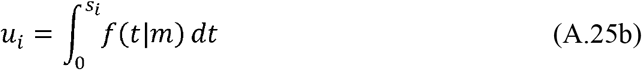

which are uniformly distributed samples over [0 1]^2^, under the correct model. Here, Θ(*i*) represents the corresponding permutated number for i. 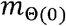 is equal to −∞ and 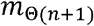 is equal to +∞.

**Proof:** We can write the joint probability distribution of the event time and mark by

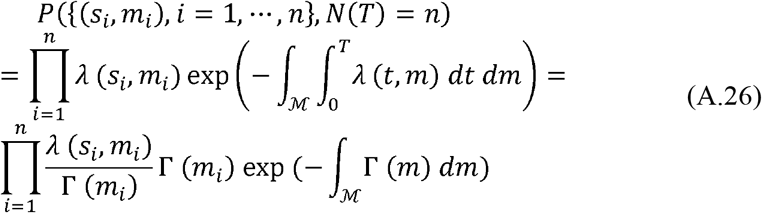

We can replace the exponential term in the Equation (A.26) using different samples’ mark information

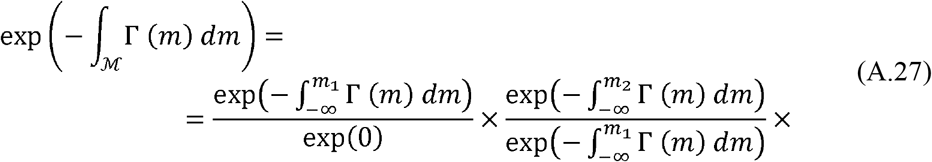

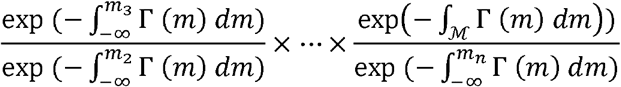

where, marks are sorted from the lowest to largest one – a lower index corresponds to a mark with a lower value. Note that the above equality is valid for any sequence of *m*_*i*_ where *i* is defined by a permutation of [1 *n*]. Thus, we have

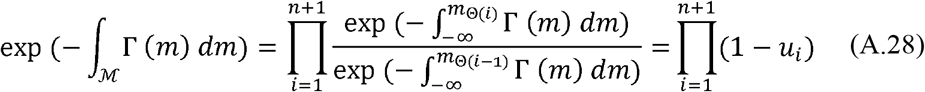

where, Θ(*i*) represents the corresponding permutated number for *i*. 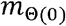 is equal to −∞ and 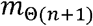 is equal to +∞. In Equation (A.28), we assume *m*_Θ(*i*)_ is larger than 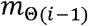, otherwise we swap the nominator and denominator – check Equation (A.25a).

Now, we can use the change of variable theorem to prove that *u*_*i*_ and *v*_*i*_ are uniformly distributed in [0 1]^2^. Partial derivative of *u*_*i*_ with respect 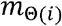 is defined by

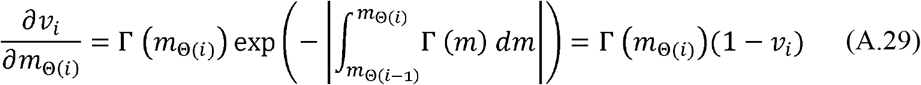

and partial derivative of *u*_*i*_ with respect *s*_*i*_ is defined by

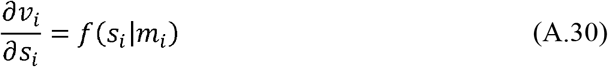

By replacing these in the joint probability distribution of (*s*_*i*_,*m*_*i*_) defined in Equation (A.26), we get the following probability distribution

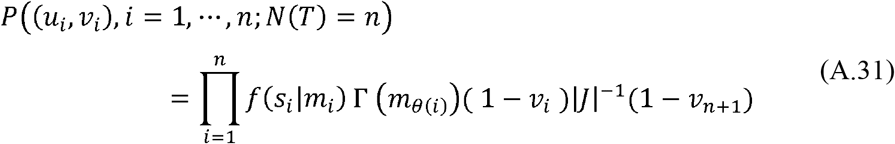

where, |*J*| is the Jacobian of transformation defined by

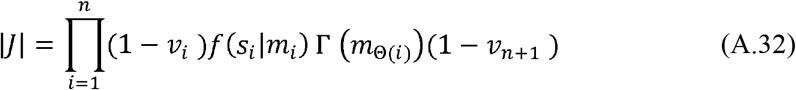

where, for the last term, we take the derivative with respect 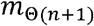, which we assume it goes to +∞. To calculate the derivative, we leverage from the fact that *J* is an upper – or lower – triangular matrix. Note that we can use the row operation to change the order of Θ(*i*) to *i* and this will only change the sign of the determinant in Equation (A.32). For the transformation defined in Equation (A.31), the sign of the determinant is not important given we only need the absolute value of the determinant. The right side of Equation (A.31) becomes equal to 1,

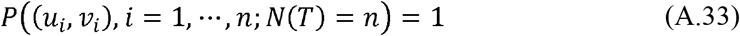

which suggests *u*_*i*_ and *v*_*i*_ are uniformly distributed samples over the space [0 1]^2^

### Corollary 3

The *v*_*i*_ samples generated by Equation (A.26a) given the ground conditional intensity defined in Equation (A.24) are uniformly distributed over the range 0 to 1 independent of *v* samples.

Here, we build a point process over the mark space, *m*; thus, if the marks are sorted in ascending order, under the time-rescaling theorem [5], *v*_*i*_s are independent and uniformly distributed random variables. The result will be like Corollary 2.

## Appendix C. Uniformity Tests

Here, we describe different terms used in the Discrepancy and Ripley Static tests.

## Appendix C.1. Discrepancy Test

In this appendix, we provide *U*_1_ and *U*_2_ terms’ definitions used in the step 4 of discrepancy test. In this test, we calculate the following equation after generating data samples by running IRCM or MDCI transformations.

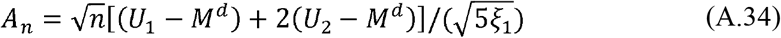

In this equation for symmetric discrepancy *M* = 4/3, 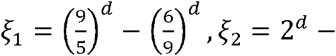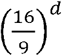 and *U*_1_, *U*_2_ dare defined by

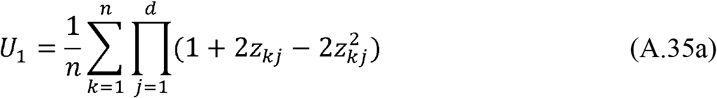

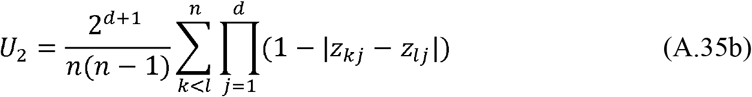

where *z*_*kj*_ are uniform samples that *k* and *j* refer to *k*^*th*^ sample and *j*^*th*^ dimension of data. For example, for simulation data *z*_*k*1_ = *u*_*k*_, *k* = 1, ‥, *n* and *z*_*k*2_ = *v*_*k*_, *k* = 1,‥,*n*.

## Appendix C.2. Ripley Statistics test

The ∑ used in step 4 of Ripley statistics method is defined by

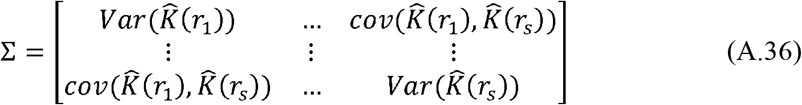

where, *r*_*i*_ ∈ *r*, *r* = {*r*_1_,…,*r*_*S*_} indicates the neighborhood radius of each sample point. The variance, 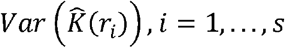 and covariance, 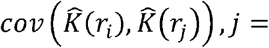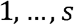 terms are defined by

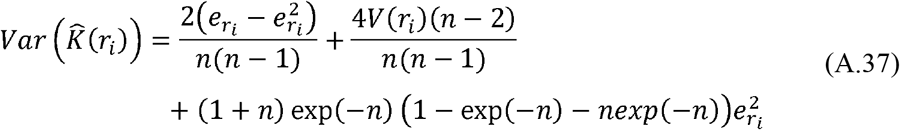

where, *n* is the number of samples and

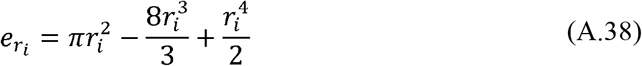

and

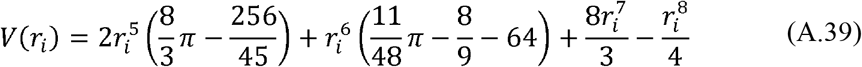

and for the covariance term

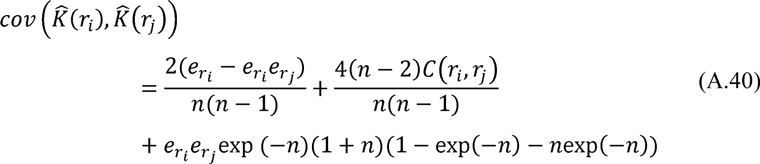

where,

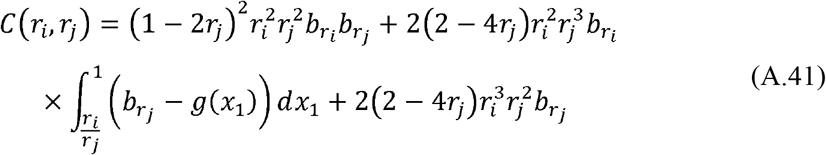

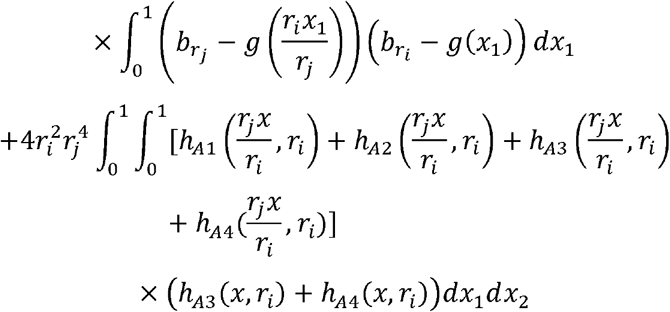

where, *x* = [*x*_1]_ *x*_2_]^*T*^ is the sample point and

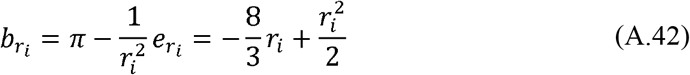

Also *g*(.) and *h*_*Ak*_(.) *k* = 1,…,4 in Equation (A.41) are defined by

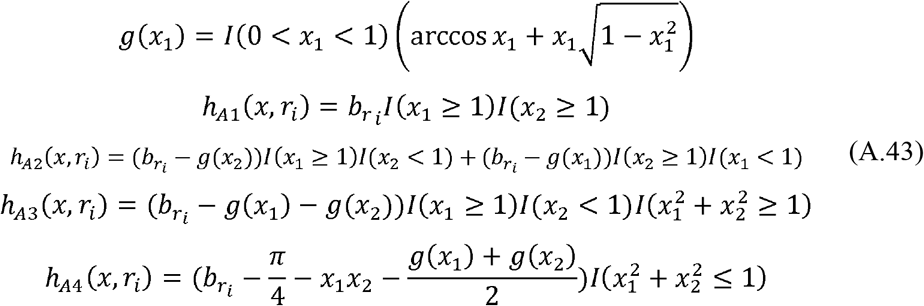

